# Human gut metagenomic mining reveals an untapped source of peptide antibiotics

**DOI:** 10.1101/2023.08.31.555711

**Authors:** Marcelo D. T. Torres, Erin Brooks, Angela Cesaro, Hila Sberro, Cosmos Nicolaou, Ami S. Bhatt, Cesar de la Fuente-Nunez

**Affiliations:** Machine Biology Group, Departments of Psychiatry and Microbiology, Institute for Biomedical Informatics, Institute for Translational Medicine and Therapeutics, Perelman School of Medicine, University of Pennsylvania, Philadelphia, Pennsylvania 19104, United States of America; Departments of Bioengineering and Chemical and Biomolecular Engineering, School of Engineering and Applied Science, University of Pennsylvania, Philadelphia, Pennsylvania 19104, United States of America; Penn Institute for Computational Science, University of Pennsylvania, Philadelphia, Pennsylvania 19104, United States of America; Department of Medicine (Hematology; Blood and Marrow Transplantation), Stanford University, Stanford, CA, United States of America; Department of Genetics, Stanford University, Stanford, CA, United States of America

## Abstract

Drug-resistant bacteria are outpacing traditional antibiotic discovery efforts. Here, we computationally mined 444,054 families of putative small proteins from 1,773 human gut metagenomes, identifying 323 peptide antibiotics encoded in small open reading frames (smORFs). To test our computational predictions, 78 peptides were synthesized and screened for antimicrobial activity *in vitro*, with 59% displaying activity against either pathogens or commensals. Since these peptides were unique compared to previously reported antimicrobial peptides, we termed them <u>s</u>mORF-encoded peptides (SEPs). SEPs killed bacteria by targeting their membrane, synergized with each other, and modulated gut commensals, indicating that they may play a role in reconfiguring microbiome communities in addition to counteracting pathogens. The lead candidates were anti-infective in both murine skin abscess and deep thigh infection models. Notably, prevotellin-2 from *Prevotella copri* presented activity comparable to the commonly used antibiotic polymyxin B. We report the discovery of hundreds of peptide sequences in the human gut.

## Introduction

A pressing threat in healthcare is the rapid emergence of antibiotic resistance in bacteria, which has often outpaced new drug discovery^1–3^. This has led to a dwindling supply of therapeutics to target multi-drug resistant pathogens, which are among the leading causes of nosocomial infections. Short peptides are a promising class of antimicrobial agents capable of addressing this threat^3^. Their short length of <50 amino acid residues, vast sequence space, and nonspecific mechanisms of action make them potential drug candidates^4^.

Antimicrobial peptides (AMPs) can be encoded by unicellular and multicellular organisms; in metazoans, for example, AMPs are an ancient form of host defense^4^. In addition to directly targeting bacteria, certain ‘host-derived’ peptides from metazoans have immunostimulatory properties that boost their potency and could be similarly leveraged by synthetic AMPs^5^. Interestingly, documented resistance to AMPs derived from prokaryotes or eukaryotes is exceedingly rare^2,3^. In those few cases where resistance has been documented, cross resistance to other AMPs ^3^ has not been demonstrated.

The physiochemical features of AMPs and the low rate of resistance to them have resulted in their use in clinical medicine. For example, AMPs that are widely used include bacitracin, colistin, and polymyxin B. Bacitracin, produced by *Bacillus licheniformis* bacteria, targets Gram-positive bacteria by interfering with cell wall and peptidoglycan synthesis. It is used to treat eye and skin infections. Polymyxin E (colistin), a last resort antibiotic produced by *Paenibacillus polymyxia* variant *collistinus*, has activity against Gram-negative bacteria, working by displacing bivalent cations as counter ions of lipopolysaccharides that form the bacterial membrane. Colistin is used mostly to treat pneumonia and biofilms in cystic fibrosis patients. Polymyxin B, produced by *Paenibacillus polymyxia,* an antibiotic used to treat topical infections and gut decontamination^6^, functions by disrupting the outer membrane of Gram-negative bacteria. Thus, AMPs have potential for clinical utility; however, few other AMPs are commercially available. This is because identifying AMP candidates has been difficult. Current methods used to discover new molecules rely primarily on prospection of naturally occurring organisms, which is slow and upredictable as it relies on trial-and-error experimentation. Additional engineering of AMPs has leveraged heuristics whereby amino acid residues substitutions are introduced^7–11^, guided by alterations in physicochemical feature determinants known to contribute to membrane targeting and, ultimately, antimicrobial activity^4^. Recently, advances in machine learning^12,13^ and methods^14^ such as generic algorithms^15^ and pattern recognition algorithms^16,17^ have yielded improved peptides. However, only a few efforts have focused on mining proteomes^18,19^ and metagenomes^18^. The human microbiome, for example, is one such space that offers significant promise in antimicrobial discovery but remains underexplored. For one, it has long been known that healthy members of the human microbiota can suppress the growth of pathogens. In addition, given the intense competition required to carve out a niche in this space, there is good reason to believe that this pool of microbial peptides should be significantly enriched for candidates with antimicrobial activity. This niche has been overlooked in recent years owing to the computational challenge of annotating proteins <50 amino acid residues, the length most common among AMPs.

Recently, we overcame this limitation and found that the human gut metagenome encodes hundreds of thousands of small open reading frames (smORFs)^18^, of which only a tiny fraction has been functionally characterized. These represent a vast untapped source of unexplored peptide sequences with potential antimicrobial activity. To narrow down the extensive list of candidate smORFs to a list that is tractable for antimicrobial activity testing, we used existing computational algorithms to predict the likelihood that a given peptide sequence has antimicrobial activity. We then chemically synthesized 78 of these peptides and tested them against high priority pathogens and commensals *in vitro*, finding 46 active peptides (59% hit rate). Five lead candidates from different sources, which had high activity against pathogens but limited or no activity against commensals, were tested and showed activity in preclinical infection animal models. Taken together, we leveraged our computational and experimental screening platform to mine a recently expanded microbiome microprotein database for candidate peptides with antimicrobial activity and identified highly potent and specific novel sequences (**Figure 1**).

**Figure 1.**
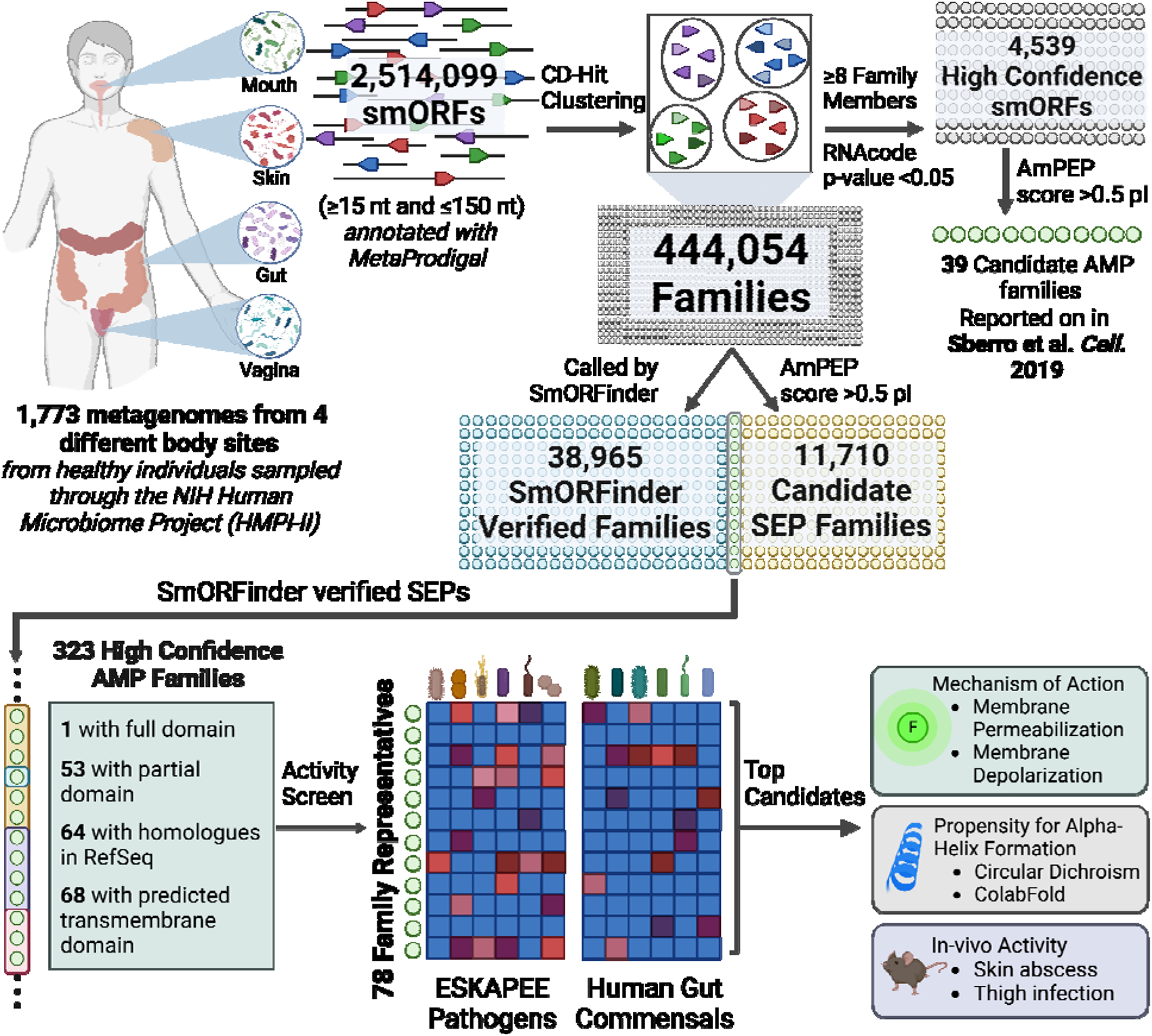
Schematic of the computational-experimental platform for the discovery of SEP from smORFs. Metagenomes from four distinct body sites were analyzed to identify open reading frames (ORFs) containing more than 15 base pairs, using the MetaProdigal tool. Subsequently, small ORFs (≤150 bp) were filtered out, and the encoded proteins were grouped into families, resulting in a total of 444,054 families. To further narrow down the selection, representatives of each family underwent analysis with SmORFinder, and the results were ranked using AmPEP to identify peptides with antimicrobial potential. The sequences that were identified by both SmORFinder and ranked as antimicrobials by AmPEP were considered as high-confidence families. These families were then subjected to further filtering based on specific criteria, as outlined in the **Inclusion and Exclusion Criteria for Selecting Peptides for Activity Testing** section. The selected high-confidence families were subsequently tested against a range of pathogen and commensal bacterial strains. Promising candidates were further investigated through systematic characterization, including conformational studies, mechanism of action elucidation, assessment of synergistic interactions, and evaluation in pre-clinical mouse models. The figure was created with BioRender.com.

### In silico identification of candidate AMPs from the human microbiome

We first developed a discovery pipeline in which promising antimicrobial candidates were identified *in silico* from smORFs annotated in human gut metagenomes. We drew on a list of 444,054 families of putative small proteins (referred to as the 444k set) that were previously predicted using a comparative genomic workflow.^7^ Briefly, MetaProdigal^8^ was used to annotate all open reading frames (ORFs), as short as 15 base pairs, on 128,368,337 contigs spanning more than 180 billion base pairs of sequenced DNA from 1,773 metagenomes from 263 healthy individuals sampled from four different major body sites. We filtered out ORFs that encode for proteins that are >50 amino acid residues in length and then clustered the remaining proteins based on sequence and length similarity using CD-Hit,^9^ resulting in the 444k set. Whereas before we focused our analysis on a subset of the 4k families with signatures of evolutionary conservation, in this study we mined the entire 444k set for candidate peptides with antimicrobial properties. We reasoned that those may be rapidly evolving or encoded by rare members of the microbiome, in which case they might not be similar to other smORFs gene products in our data set. Using AmPEP, a random forest classifier that predicts whether a given query sequence is likely to have antimicrobial activity or not, we found 11,710 smORFs families that have an AmPEP score of ≥0.5 which corresponds to a score that indicates a given amino acid sequence is likely to be an AMP.^11^ To determine which of these smORFs are genuine protein-encoding genes, we determined which ones were also identified by SmORFinder,^6^ a tool that combines profile hidden Markov models of each smORF family and deep learning models to predict smORF families that are likely to be valid. This reduced the list to 323 smORF-encoded candidate peptides with predicted antimicrobial activity (**Data S1**), from which we selected 78 to synthetize and screen for biological activity. Selection of the final 78 peptides was based on four criteria: (i) sequences with a high AmPEP score were preferentially selected, (ii) representation of the family of origin of the peptide (**Data S1**); *e.g.*, clustered families with more members were preferentially selected over families with just one or a few candidate sequences; (iii) amino acid residues composition that would enable an effective chemical synthesis, *e.g.*, sequences with motifs composed of amino acid residues hindered side chain that would lead to low yield or many cysteine residues indicating constrained and complex secondary structures were filtered out; and (iv) sequences without hydrophobic clusters were selected to avoid aggregation that would interfere with our subsequent screening effort.

### Physicochemical features reveal a new class of candidate peptide antibiotics

Physicochemical descriptors play a pivotal role in elucidating the activity of AMPs. These descriptors provide valuable insights into the structural and molecular properties of AMPs, allowing for a deeper understanding of their interactions with microbial membranes, their ability to penetrate cell membranes, and their stability in various biological environments. By quantitatively characterizing the physicochemical features of AMPs, it is possible to perform informed predictions about their antimicrobial activity, aiding in the design and optimization of more effective peptides. Thus, to assess the physicochemical features of all predicted (323 families) candidate smORF-encoded AMPs, we analyzed the amino acid residues composition (**Figures 2a and S1**) and known physicochemical determinants of antimicrobial activity (**Figures 2b-c and S2**). We calculated their main physicochemical features using the Database of Antimicrobial Activity and Structure of Peptides (DBAASP) server^20^ (**Figures 2b-c and S2**). For comparison purposes, we selected AMPs listed in DBAASP and encrypted peptides (EPs) found in the human proteome with predicted antimicrobial activity^21^, which are two classes of peptide antibiotics with different physicochemical features, amino acid composition, and mode of action. Despite the fact that AmPEP predicts antimicrobial activity based on features found in known AMPs within its training set, the 323 SEP candidates displayed a higher content of negatively charged residues (aspartic and glutamic acids) than known AMPs and EPs (**Figures 2a and S1a-d**). This impacted their overall net charge, which is lower than most described AMPs and EPs (**Figure 2b**). The candidate SEPs also had a higher content of aliphatic and hydrophobic amino acid residues, in general, than hydrophilic ones (polar uncharged, acidic, and basic) compared to known AMPs and EPs (**Figures 2a and S1b-d**). Despite their high content of aliphatic and aromatic residues (**Figure S1b-d**), the low hydrophobic moment values (**Figure S2a**) summed to the high content of polar uncharged residues, such as methionine and asparagine, and to negatively charged residues make SEP candidates substantially less hydrophobic than previously described AMPs and EPs (**Figures 2c, S1 and S2**). The averaged lower hydrophobicity and net charge of the candidate SEPs directly influences their higher tendency to disordered conformation (**Figure S2b**), lower isoelectric point (**Figure S2c**), and lower amphiphilicity (**Figure S2d**) compared to already reported AMPs and EPs. It is worth noting that the high content of methionine residues (**Figure 2a**) was expected in our peptide sequences since its associated codon AUG is the most common translation initiator^22^. These physicochemical data also reveal that these peptides do not rely on amphipathic structures with aliphatic and positively charged residues but instead consist of methionine-rich, slightly less hydrophobic sequences with an increased tendency to be disordered. Altogether, these data suggest that the peptides identified in this work constitute a new class of sequences, separate from known AMPs and EPs.

**Figure 2.**
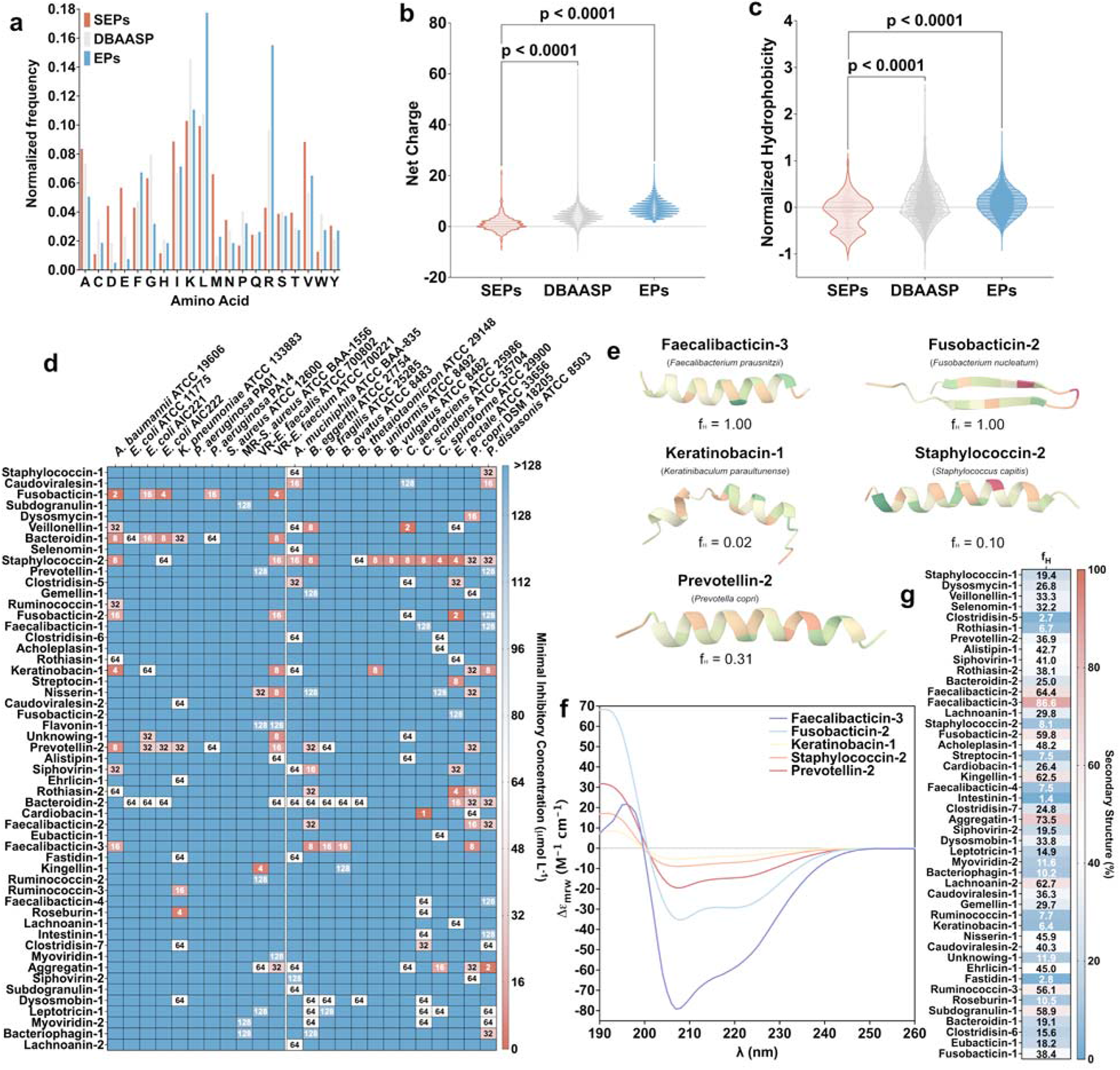
Sequence-related features, antimicrobial activity, and structure analysis of SEPs. **(a)** Amino acid frequency was calculated from candidate SEPs, from EPs predicted in the human proteome and from AMPs the DBAASP database. SEPs had overrepresentation of acidic residues (aspartic acid – D and glutamic acid – E), polar uncharged residues (methionine – M, and asparagine – N), and lower content of leucine (L) residues. Among the most relevant physicochemical features that are known to influence biological activities of peptides (see also **Figure S1**), **(b)** SEPs have lower net positive charge and **(c)** normalized hydrophobicity than AMPs and EPs. Thus, SEPs are not amphipathic as other classes of AMPs, instead they are slightly less hydrophobic sequences with higher tendency to be disordered (see also **Figure S2** and **Table S1**). **(d)** Antimicrobial activity of the tested SEPs. Briefly, a 10^6^ bacterial cells load was exposed to serially diluted SEPs (1-128 μmol L^-1^) in 96-well plates and incubated at 37LJ°C. Twenty hours after the beginning of the experiment, each condition was analyzed in a microplate reader at 600LJnm to check for inhibition of bacterial growth compared with the untreated controls. The results are presented as a heat map of antimicrobial activities (μmol L^-1^) against 11 pathogenic and 8 gut commensal bacterial strains. Assays were performed in three independent replicates. **(e)** ColabFold was used to generate structural predictions using default parameters. Three-dimensional ribbon structures of the resulting PDB files were generated using Mol*3D Viewer. **(f)** Circular dichroism spectra of the SEPs in helical inducer medium to assess the tendency of SEPs to the most common structure adopted by antimicrobial peptides. Five of the SEPs present significant helical content (faecalibacticin-3, fusbacticin-2, keratinobacin-1, staphylococcin-2, and prevotellin-2) expressed in **(g)** helical fraction (f_H_) values in a heat map, where the higher f_H_ values are presented in red and the lowest in blue. The activity did not correlate with the antimicrobial activity, once again reinforcing the independency of this class of peptides from amphipathic and balanced hydrophobic/cationic residues sequences (see also **Figure S3a-b**).

### Antimicrobial activity of SEPs

To assess the potential antimicrobial activity of the candidate SEPs, we chemically synthesized 78 sequences (**Table S1**) and tested them against 11 clinically relevant pathogenic strains (*Acinetobacter baumannii*, three *Escherichia coli* strains including a colistin-resistant strain, *Klebsiella pneumoniae*, two *Pseudomonas aeruginosa* strains, two *Staphylococcus aureus* strains including a methicillin-resistant strain, vancomycin-resistant *Enterococcus faecalis*, and vancomycin-resistant *Enterococcus faecium*), several of which are considered ESKAPEE pathogens (*E. faecium*, *S. aureus*, *K. pneumoniae*, *A. baumannii*, *P. aeruginosa*, *Enterobacter spp*., and *E. coli*), and thus the most threatening bacterial pathogens in our society according to the World Health Organization (**Figure 2d**). Briefly, a gradient of concentrations ranging from 1 to 128 μmol L^-1^ of the peptides were exposed to 10^6^ cells mL^-1^ of each one of the strains in the same medium (Luria-Bertani) and in sterile 96-well plates. Those plates were subsequently incubated for 24 h at 37 °C. After this period, the optical density at 600 nm (OD_600_) was measured in an absorbance reader and the minimal inhibitory concentrations (MIC) values were determined as the concentrations where the peptide completely inhibits the growth of the bacteria (MIC_100_). This initial screen yielded 25 SEPs (32% of the synthesized SEPs) that completely sterilized bacterial loads of at least one of the pathogens tested. Interestingly, the only strain that was not targeted by any of the SEPs was the Gram-positive bacterium *S. aureus*.

Next, since SEPs were identified from human-associated metagenomes across several different body sites, including the gut, we screened them against 13 of the most abundant members of the human gut microbiota to assess whether they were able to target gut commensals^23^. The following bacteria were tested: *Akkermansia muciniphila*, *Bacteroides eggerthi*, *Bacteroides fragilis*, *Bacteroides ovatus*, *Bacteroides thetaiotaomicron*, *Bacteroides uniformis*, *Bacteroides vulgatus* (*Phocaeicola vulgatus*), *Colinsella aerofaciens*, *Clostridium scindens*, *Clostridium spiroforme*, *Eubacterium rectale*, *Prevotella copri*, and *Parabacteroides distasonis* belonging to four different phyla (Verrucomicrobia, Bacteroidetes, Actinobacteria, and Firmicutes). The experiments were performed as described for the pathogenic strains but under anaerobic conditions and in brain heart infusion medium. Classical antimicrobial peptides (AMPs) usually do not target microbiome strains^24^, however we previously found that EPs were able to kill commensals. Our screen yielded 39 SEPs with low micromolar antimicrobial activity against at least one of the gut commensals tested (50% hit rate). We also found that all gut microbiome strains tested were susceptible to at least one SEP. In total, 46 of the 78 synthesized SEPs (59%) had antimicrobial activity against at least one pathogen or commensal.

### Identification of organisms encoding SEPs

While candidate AMPs are often tested for activity against pathogens, relatively little is known about AMP activity against commensals of the human microbiome. Given the high number of SEPs that displayed antimicrobial activity against commensals, we sought to learn more about the potential source organism for each of these. We evaluated contigs containing each of the original 323 predicted SEPs from our initial list and ran a BLAST search against NCBI’s nucleotide database. We chose the organism of the top hit for every query as the putative source organism. Overall, of the 285 SEPs for which the genus of the source organism could be predicted, 69 different genera were represented. Consistent with our hypothesis that SEPs would be encoded by rare members of the microbiome, we found that 38 SEPs were annotated on contigs without hits or with hits to uncultured or unclassified organisms or plasmids. While many organisms on the list would be considered human commensals, including *Faecalibacterium*, *Prevotella*, *Bacteroides*, and *Lachnospiraceae* species, it was notable that several were known human pathogens or opportunistic pathogens, such as *Haemophilus parainfluenzae*, *Gemella haemolysans*, and *Escherichia coli*. Interestingly, 78 (24%) of the 323 SEPs appeared to originate from viral or phage genomes, 13 of which belonged to the list of 78 SEPs chosen for testing. In addition, one SEP appeared to be from a human contig.

We found two cases where two SEPs were encoded by the same contig: ruminococcin-1 with ruminococcin-3 (*Ruminococcus bicirculans*), and caudoviralesin-1 with caudoviralesin-2 (*Caudovirales*). However, we found that no member of either of these pairs displayed notable activity against either pathogens or commensals, suggesting that they may fill another role, either individually or in tandem. Six members of the list of 323 candidate AMPs were previously confirmed to be transcribed and translated through RNA-seq and MetaRibo-seq^25^. Therefore, it was notable that two of these six (fusobacticin-1 and bacteroidin-1) encoded by *Fusobacterium periodonticum* and *Bacteroides salanitronis*, respectively, were active with low MICs against several pathogens. Given that both SEPs were encoded by gut commensals, their expression may play a key role in colonization resistance, especially against pathogenic strains of *E. coli* and *E. faecium*.

Staphylococcin-2 showed activity against the pathogens *A. baumanii* (8 μmol L^-1^) and *E. faecium* (16 μmol L^-1^) and displayed low MICs against nearly all commensals in our panel. Interestingly, it is encoded by a contig classified to *Staphylococcus capitis,* a member of the human skin microbiome. Closer examination of the contig (likely a plasmid) revealed the SEP was encoded 704 base pairs upstream from a type I toxin-antitoxin system Fst family toxin. A cursory search of other assemblies in NCBI containing this toxin revealed that it is encoded near staphylococcin-2 in several, but not all assemblies examined.

Another interesting SEP was Bacteroidin-2, which showed moderate activity (MIC values ranging from 16 – 64 μmol L^-1^) against *E. coli,* vancomycin-resistant *E. faecium* and many of the commensals in our panel. This SEP was encoded by a contig classified to *Bacteroides cellulosilyticus*, a common member of the human gut microbiome known to ferment complex carbohydrates, including cellulose. It could be possible that *B*. *cellulosilyticus* uses bacteroidin-2 to maintain a niche within a healthy gut.

### Secondary structure of the active SEPs

The secondary structure of peptides dictates their antimicrobial and other biological activities. Since the synthesized SEPs were short (13-44 amino acid residues, with most containing around 25 residues), they tended to be disordered in hydrophilic environments, such as water and buffers. To generate a first approximation of possible structure, we ran the 323 candidate SEP sequences through ColabFold using default parameters. ColabFold predicted all but two of the peptides to contain an α-helical domain. The two exceptions, SEPs fusobacticin-2 and roseburin-1, contained a β-like structure or were completely disordered, respectively. Although AMPs with high activity are often found to be α-helical, ColabFold predicted α-helical structures for both SEPs found to be active and those found to be inactive. Some of the predicted structures were interesting in that two distinct domains were predicted; an α-helical domain joined with a disordered region, which was the case for bacteroidin-2, clostridisin-5, and alistipin-1. One of the SEPs, staphylococcin-1, was notable for its kinked predicted α-helical structure. Five of the most active SEPs ColabFold structures are shown in **Figures 2e**. Given that AlphaFold is known to predict high confidence α-helical structures for even spurious small proteins (<100 amino acid residues)^26^, we probed the secondary structure tendencies of the peptides through Circular Dichroism in trifluoroethanol (TFE) and water mixtures (3:2, v:v). This solvent is a known α-helical inducer by dehydrating the amide groups that are part of the backbone of the peptide sequence and favoring intramolecular hydrogen bonds that lead the peptide to helical conformation^27,28^.

First, we selected the active SEPs and diluted the peptides in the TFE/water mixture at a fixed concentration (50 μmol L^-1^), obtaining the circular dichroism spectra for wavelengths ranging from 260 to 190 nm (**Figures 2f and S3a**). To obtain the secondary structure fractions we used the Beta Structure Selection (BeStSel) server^29^ (**Figures 2g and S3b**). As expected, the ColabFold predicted secondary structures poorly correlated with the secondary structure fractions obtained experimentally (**Figure S3b**). Most of the SEPs presented intermediary and low values of α-helical content in the conditions tested, and only eight of them presented helical fraction (f_H_) values higher than 50%. These results support the conclusion that SEPs tend toward a disordered conformation (**Figure S3b**), and that this does not adversely affect their antimicrobial activity.

### Synergy between SEPs

Next, we wondered whether SEPs from the same biogeographical area of the body could synergize to target bacteria (**Figure 3a**). Checkerboard assays^21,30^ were performed with pairs of SEPs derived from the same body sites: i.e., the tongue dorsum, supragingival plaque, and stool. To quantify the interactions between the SEP pairs, we extracted their fractional inhibitory concentration index (FICI)^31^. Remarkably, almost all SEP pairs tested experimentally displayed synergistic or additive interactions against the Gram-negative pathogen *A. baumannii*; the concentrations needed ranged from low micromolar to nanomolar concentrations *in vitro*, doses comparable to the most potent AMPs^10,32,33^ and EPs^21,34^. One of the peptide pairs (veilonellin-1 and rothiasin-2) from the metagenome of bacteria from the tongue dorsum presented the most significant synergistic interaction with a FICI of 0.25.

**Figure 3.**
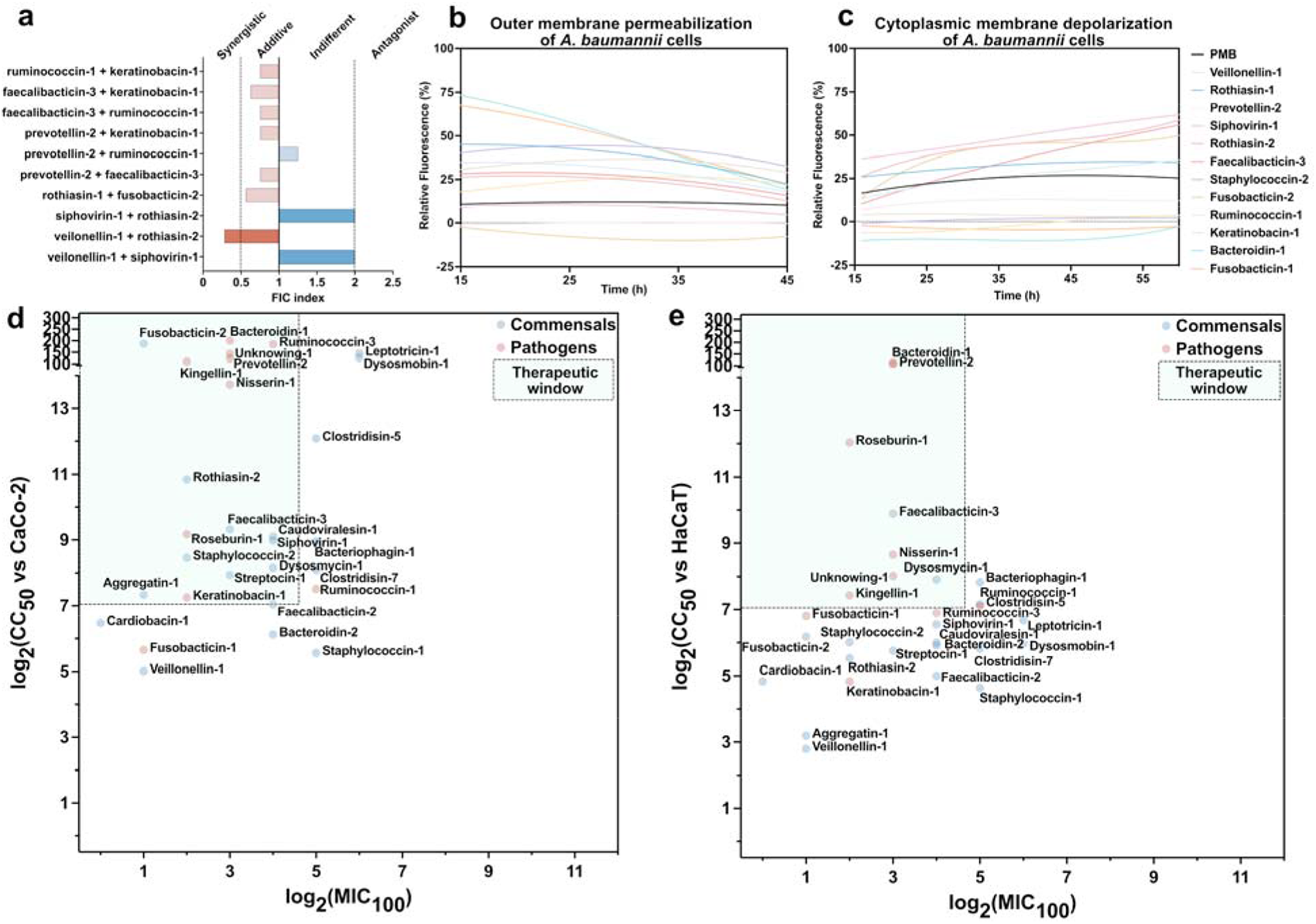
Synergy, mechanism of action, and cytotoxicity of SEPs. **(a)** The synergistic interaction between pairs of SEPs from the same biogeography (tongue dorsum, supragingival plaque, and stool) was assessed by checkerboard assays with 2-fold serial dilutions starting at 2×MIC to MIC/32. The histogram shows the fractional inhibitory indexes (FICI) values obtained for each pair of SEPs, where dark red represents synergistic interactions, light red indicates additive interactions, and blue shows indifferent interactions. Most of the pairs of SEPs presented synergistic or additive interactions. To assess whether SEPs act on the bacterial membrane, all active SEPs against each of the pathogenic strains were tested in outer membrane permeabilization and cytoplasmic membrane depolarization assays. In general, SEPs presented low permeabilization of the outer membrane effect, as shown in **(c)** the relative fluorescence measurements of SEPs on *A. baumannii* cell membranes (see also **Figure S4**). SEPs showed high depolarization properties as shown in **(d)** the relative fluorescence measurements of SEPs on vancomycin-resistant *E. faecium* cytoplasmic membranes (see also **Figure S5**). The relative fluorescence was calculated with a non-linear fitting using as baseline the untreated control (buffer + bacteria + fluorescent dye) as described in the methods section. **(e)** The correlation between cytotoxicity on human colorectal adenocarcinoma cells (Caco-2) and antimicrobial activity is shown in a scatter plot where the cytotoxicity is represented by the CC_50_ values (cytotoxic concentrations causing 50% cell death) and MIC (minimal inhibitory concentration for complete bacterial killing). CC_50_ values have been predicted by interpolating the dose-response with a non-linear regression curve. The green area represents the therapeutic window where those peptides could be safely used with no toxic effect to eukaryotic cells (see also **Figure S6**).

### Mechanism of action of SEPs

To investigate how SEPs exert their effects on bacterial cells, we conducted fluorescence assays to determine if these peptides act by targeting the membrane. Firstly, we identified all antimicrobial hits among the SEPs (**Figure 2d**). Subsequently, we evaluated the capacity of these peptides (at their MIC value) to disrupt (**Figure 3b and S4**) and depolarize (**Figure 3c and S5**) the bacterial the outer and cytoplasmic membrane, respectively. To assess whether SEPs can permeabilize the outer membrane of Gram-negative bacteria, we performed 1-(N-phenylamino)naphthalene (NPN) assays. NPN is a lipophilic dye that emits fluorescence in lipid-rich environments like bacterial outer membranes. If the bacterial outer membrane is damaged, NPN can permeate and increase its fluorescence (**Figure 3b**). The following Gram-negative strains were exposed to SEPs in the presence of NPN: *A. baumannii* ATCC 19606 (**Figure S4a**), *E. coli* ATCC 11775 (**Figure S4b**), *E. coli* AIC221 (**Figure S4c**), *E. coli* AIC222 (**Figure S4d**), *K. pneumoniae* ATCC 13883 (**Figure S4e**), and *P. aeruginosa* PA14 (**Figure S4f**). All strains were permeabilized by the SEPs, except for *E. coli* ATCC 11775, which was permeabilized by all SEPs except for SEP bacteroidin-2 (this peptide was only active in terms of antimicrobial activity against *E. coli* strains) (**Figure 2d**). Similarly, *P. aeruginosa* PA14 was not permeabilized by prevotellin-2, a broad-spectrum SEP that did permeabilize *A. baumannii* ATCC 19606, *E. coli* strains AIC221 and AIC222, and *K. pneumoniae* ATCC 13883. The peptide antibiotic polymyxin B was used as a positive control in our studies^21^. In summary, SEPs did not permeabilize the outer membrane of bacteria to the level previously reported for AMPs^7,32^ or EPs^21^, indicating that the mechanism of action of SEPs might be independent of outer membrane permeabilization.

Next, we used 3,3′-dipropylthiadicarbocyanine iodide (DiSC_3_-5), a fluorophore used to assess whether a compound depolarizes the bacterial cytoplasmic membrane. If there are imbalances in the transmembrane potential of the cytoplasmic membrane, the fluorophore migrates to the extracellular environment, producing fluorescence. Out of all 48 conditions tested, in 40 occasions SEPs tested depolarized the cytoplasmic membrane of bacteria more substantially than groups treated with polymyxin B, a control which displays certain level of depolarization^21^ (**Figure S5a-i**). Overall, these data suggest that SEPs operate preferentially via depolarizing the cytoplasmic membrane as opposed to permeabilizing the outer membrane, revealing a mechanism that is distinct from that of conventional AMPs^7,32^ and EPs^21^, which tend to target the outer membrane.

### Cytotoxicity assays

To assess the potential toxicity of SEPs towards mammalian cells, we tested the 29 SEPs with higher antimicrobial activity. The peptides were exposed to human colorectal adenocarcinoma cells (Caco-2) and immortalized human keratinocytes (HaCaT), which serve as models of intestine and skin epithelia, respectively^35–37^. In the case of Caco-2 cells, most peptides showed CC_50_ values (*i.e.*, cytotoxic concentration that leads to 50% cell death) higher than 32 μmol L^-1^ (**Figure 3d and S6a**). Nearly all sequences active against bacterial pathogens at low MIC values did not display toxic effects when tested on Caco-2 cells, except for peptides staphylococcin-1, veillonellin-1, and fusobacticin-1, which were toxic at 32, 32, and 64 μmol L^-1^, respectively (**Figure S6a**). Interestingly, the predicted CC_50_ values of SEPs when exposed to human keratinocytes were lower than those obtained for colorectal cells (**Figure 3e and S6b**). Particularly, peptides staphylococcin-1, veillonellin-1, and aggregatin-1 displayed toxic effects when tested at 16, 8 and 4 μmol L^-1^, respectively, towards HaCaT cells. The higher cytotoxic profile found for HaCaT rather than Caco-2 cells might be explained by the differences in lipid membrane composition. Nontumorigenic cells, such as HaCaT, present a lower content of negatively charged phospholipids exposed to the outer side of the membrane with respect to tumoral-originated cells such as Caco-2. The presence of negatively charged and polar uncharged residues (**Figure 2a**) in the SEPs’ sequences hinders their electrostatic interactions with Caco-2 cell membranes ^38–40^. Such interactions are known to play a crucial role as primary interactions between peptides and lipid bilayer and promote their approximation to the interface membrane-extracellular environment ^4^. Keratinocytes were generally more susceptible than Caco-2 cells at relatively low concentrations of SEPs (4-8 μmol L^-1^), although those values were higher than the values obtained in the antimicrobial assays (MICs). Therefore, to prioritize which SEPs to test in animal models, we delineated the therapeutic window of each sequence (**Figure 3d-e**), to ensure that the cytotoxic concentration of each peptide against both cell lines tested was at least 2-fold lower than their antimicrobial activity, *i.e.*, 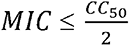 .

### Anti-infective activity of SEPs in two different preclinical animal infection models

To test if the lead SEPs retained their antimicrobial potency in complex living systems, we tested them in two mouse models, namely in mouse skin abscess^34,41,42^ and deep thigh infection^19,21^ models (**Figure 4a**). In both models, we used the pathogen *A. baumannii*, which causes infections in the blood, urinary tract, lungs, and topical wounds and is one the major causes of mortality in hospitalized patients^43^. Five lead SEPs displayed potent activity against *A. baumannii*: prevotellin-2 (8 μmol L^-1^), faecalibacticin-3 (16 μmol L^-1^), staphylococcin-2 (8 μmol L^-1^), fusobacticin-2 (16 μmol L^-1^), and keratinobacin-1 (4 μmol L^-1^), and thus were tested in both mouse models at their MIC value (**Figure 2d**).

**Figure 4.**
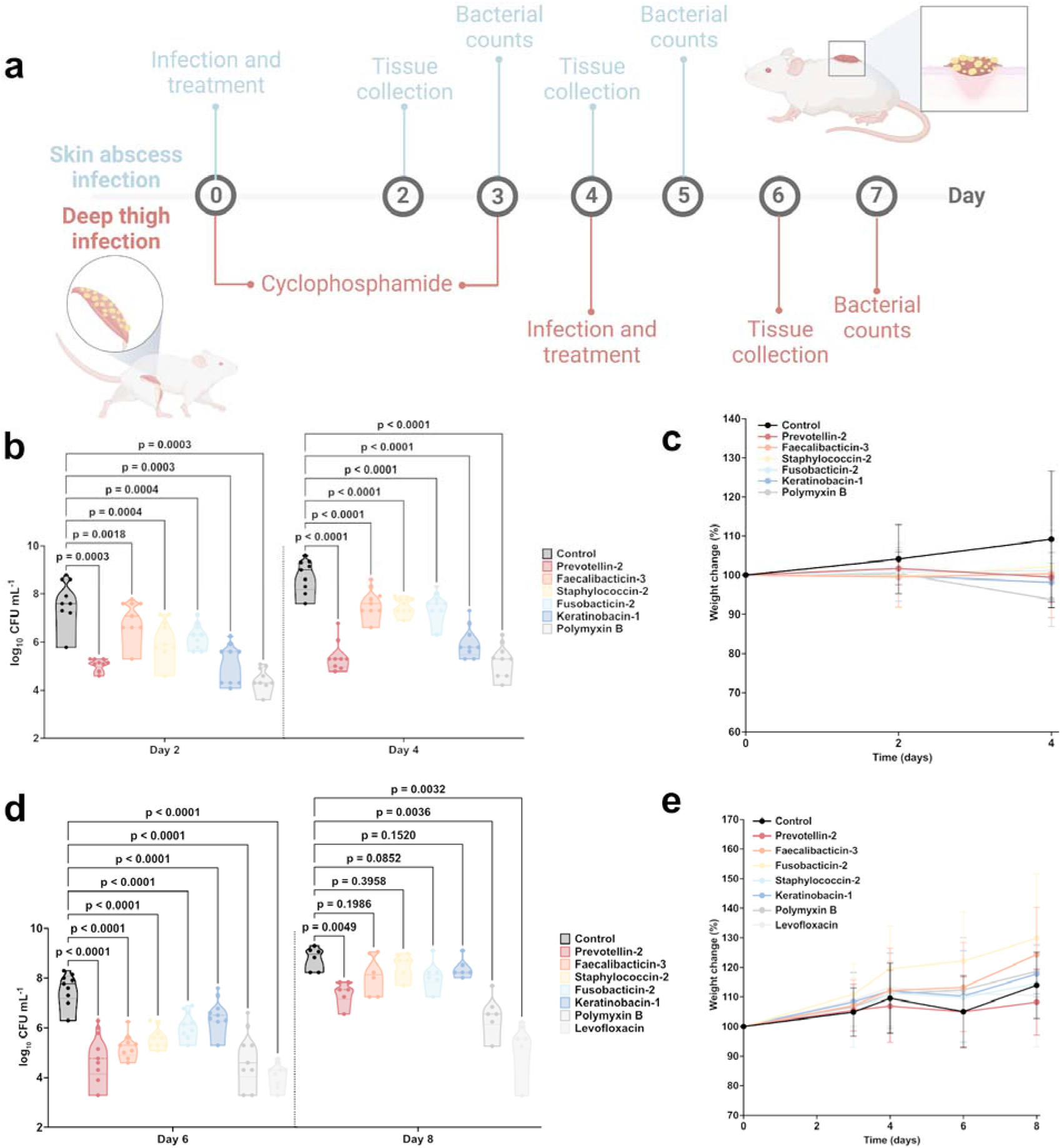
Anti-infective activity of smORF-encoded peptides (SEPs) in pre-clinical animal models. **(a)** Schematic of the skin abscess and deep thigh infection mouse models used to assess the anti-infective activity of the SEPs against *A. baumannii* cells. **(b)** In the skin abscess infection model, mice were infected with a load of *A. baumannii* and treated two hours after infection with one dose of the SEPs at their MIC. Mice were euthanized two-and four-days post infection. Each group (treated and untreated) consisted of three mice (nLJ=LJ3) and the bacterial loads used for infection of each mouse came from a different inoculum. The experiment was done in three independent replicates. All peptides had similar bacteriostatic effect two days after infection, and after four days, all the SEPs tested were significantly different than the control and the peptide prevotellin-2 presented activity comparable to the positive control (polymyxin B), reducing the infection by four orders of magnitude. **(c)** To rule out toxic effects of the peptides, mouse weight was monitored throughout the whole extent of the experiment. We considered 20% weight change as acceptable considering the duration of the experiment. **(d)** In the deep thigh infection mouse model, mice were first immunosuppressed by two rounds of treatment (24 and 72 h pre-infection) of immune system suppressor (cyclophosphamide). Subsequently, an intramuscular injection of A. baumannii was administered in the right thigh, followed by intraperitoneal administration of the peptides, to evaluate their systemic anti-infective activity two hours after infection. Six days after the start of the experiment, corresponding to two days post-infection, mice were euthanized. Each group, comprising treated and untreated mice, consisted of three individuals (n = 3), with distinct bacterial loads used for infecting each mouse, originating from different inocula. The experiment was done in three independent replicates. All peptides presented significant activity (one to two orders of magnitude reduction in bacterial counts) and the SEP prevotellin-2 had similar effect than the antibiotics used as positive controls, polymyxin B and levofloxacin, reducing three to four orders of magnitude the bacterial counts two days post-infection (day 6 of the experiment). Four days post-infection, for all treatments with SEPs, it is observed growth in bacterial counts, but prevotellin-2 and fusobacticin-2 were still as active as polymyxin B. Levofloxacin was the only treatment that led to a more than 2 orders of magnitude decrease compared to the untreated control. **(e)** During the entire 8-day period of the deep thigh infection model, mouse weight was closely monitored to eliminate the possibility of any toxic effects caused by cyclophosphamide injections, bacterial load, and the peptides. To determine statistical significance in figures **b** and **d**, one-way ANOVA followed by Dunnett’s test was employed, and the respective p-values are presented for each group. All groups were compared to the untreated control, and the violin plots display the median and upper and lower quartiles. Data in **c** and **e** are the mean plus and minus the standard deviation. Figure created with BioRender.com.

The skin abscess infection was established with a bacterial load of 10^6^ cells in 20 μL of *A. baumannii* onto a wounded area of the skin (**Figure 4a**). A single dose of each SEP at their respective MIC was delivered to the infected area. Two days post-infection, peptide prevotellin-2 markedly reduced the bacterial load by three orders of magnitude compared to the untreated control group. Its potency was comparable to the activity observed in the positive control group of mice treated with polymyxin B (**Figure 4b**). The other SEPs reduced the bacterial load by two orders of magnitude (**Figure 4b**). Four-days post infection all SEPs and polymyxin B were still preventing bacterial growth, and prevotellin-2 and polymyxin B reduced the bacterial counts by three to four orders of magnitude compared to the untreated control. All the other SEPs reduced bacterial load by two to three orders of magnitude compared to the untreated control (**Figure 4b**). These results are promising since the SEPs were administered only once and after the establishment of the abscess, highlighting their anti-infective potential. Critically, no significant changes in weight, a proxy for toxicity, were observed in our experiments (**Figure 4c**).

Next, we assessed the efficacy of the same lead SEPs (prevotellin-2, faecalibacticin-3, staphylococcin-2, fusobacticin-2, and keratinobacin-1) in a murine deep thigh infection model (**Figure 4a**). This preclinical model is widely used to assess the antibiotic potential of novel compounds. Briefly, mice were administered two rounds of cyclophosphamide treatment for immunosuppression before the intramuscular infection with 10^6^ cells in 100 μL of the bacterial pathogen *A. baumannii*. A single dose of each SEP (at their MIC) was delivered intraperitoneally (**Figure 4a**). Two days post-treatment, prevotellin-2 and the antibiotics polymyxin B and levofloxacin (positive controls) reduced the bacterial load by three to four orders of magnitude (**Figure 4d**). All the other peptides led to a one to two orders of magnitude decrease in bacterial counts compared to the untreated control group of mice (**Figure 4d**). Four days post-treatment, the bacterial counts increased for all peptide treatment conditions and the treatment with polymyxin-B, while levofloxacin was still significantly active. In our experiments (**Figure 4e**), we did not observe any significant changes in weight, indicating that the SEPs are non-toxic. The *in vivo* results support the antibiotic properties of SEPs under physiological conditions and provide a strong basis for advancing their development as potential antimicrobial agents.

## Discussion

Here, we find that human microbiomes encode thousands of small open reading frames (smORFs) with gene products that hold great promise as novel antimicrobials. We curated a list of 323 high confidence smORF encoded peptide (SEP) families predicted to be expressed and therefore capable of playing a role in direct bacterial competition. We synthesized and tested 78 of these and found that more than half displayed antimicrobial activity against at least one pathogen or commensal. The 46 active SEPs were subjected to detailed characterization to determine their mechanism of action, secondary structure, and toxicity towards human cell lines. Interestingly, the five most promising SEPs were encoded by diverse phyla from oral, skin, and gut body sites: faecalibacticin-3 (*Faecalibacterium prausnitzii*), fusobacticin-2 (*Fusobacterium nucleatum*), keratinobacin-1 (*Keratinibaculum paraultunense*), staphylococcin-2 (*Staphylococcus capitis*), and prevotellin-2 (*Prevotella copri*). We tested these SEPs at their MIC to determine their *in vivo* anti-infective activity in skin abscess and deep thigh infection mouse models of *A. baumannii*. Our lead candidate, prevotellin-2, cleared bacterial loads at a comparable level to the current gold standard polymyxin B and without notable toxicity to the mammalian host in either infection model. Prevotellin-2, named for its producing organism, the human gut commensal *Prevotella copri*, represents, to the best of our knowledge, one of the first reports of an antimicrobial peptide produced by this species.

Upon analysis of relationships between producer and target organisms made apparent by our antimicrobial assays (**Figure 2b**), we observed three different general patterns of antagonism: (1) intra-species antagonism, (2) inter-species within body-site antagonism, and (3) inter-species body-site exclusion. The first is the least surprising given the inherent threat posed by other strains occupying the same niche in the competition for resources. Narrow spectrum AMPs have been found to be produced by members of every major phylum of bacteria as well as in some Archaea and serve a crucial role in allowing producing strains to outcompete closely related bacteria with similar metabolic needs and lifestyle. These antimicrobials include the colicins of *Escherichia coli*^44^, the pyocins of *Pseudomonas aeruginosa*^45^, the halocins of halobacteria^46^, subtilin of *Bacillus subtilis*^47^, the lantibiotics of lactic acid bacteria (LAB), and other bacteriocins. In our dataset, the most notable example in this category is the antagonism observed between prevotellin-2 and *P. copri* DSM 18205 tested in our panel of commensals. Our ability to observe further examples in this category is limited by the size of our panel.

Next, looking solely at our commensal activity panel, we observe two main categories of interspecies, broad-spectrum antagonism: within body-site antagonism and between body-site exclusion. The former, which was the most common in our dataset, included examples of varying ranges of target taxonomic specificity. Broader spectrum SEPs included several types of target class specificity. SEP faecalibacticin-3 from the gut microbiome (produced by *Faecalibacterium prausnitzii*, of the phylum Firmicutes) displayed cross-phylum antagonism targeting several specific organisms from the phylum Bacteroidetes. This pattern of phylum specific broad-spectrum antagonism is similar to the Bacteroidetes specific killing reported for bacteroidetocins encoded by certain Bacteroides species^48,49^. On the other hand, we also observe bacteroidin-2 (produced by *Bacteroides cellulosilyticus*) which targeted several different classes including nearly all Bacteroidia in our panel (except for *B. uniformis* and *B. vulgatus*), as well as *Akkermansia muciniphila*, and *Eubacterium rectale* of the Verrucomicrobiae and Clostridia classes, respectively.

Finally, we observe two examples of SEPs from other body sites targeting the gut commensals screened for sensitivity in our panel. The most striking example is staphylococcin-2 (produced by *Staphylococcus capitis*) from the skin microbiome, which displays broad activity across several phyla of gut commensals. In addition, we find it notable that fusobacticin-2 (produced by *Fusobacterium nucleatum*) from the oral microbiome has highly potent activity against the gut commensal *Eubacterium rectale*. We believe SEPs like these may play a role in shaping the niche of the producer organism by excluding non-native microbes.

Although we find SEPs encoded by human associated microbes with high levels of antimicrobial activity against both pathogens and commensals, our approach has several limitations. First, we synthesized and tested the SEPs as they were translated from the coding sequence, which caused us to miss detecting SEPs that require post-translational modifications to achieve activity^50^. However, this may be of negligible concern given that all SEPs were computationally predicted to be antimicrobial as translated. This is supported by the high percentage of SEPs (59%) that showed activity in our screen. Second, given the limitations of Prodigal and smORFinder, which we relied upon to identify and verify our list of SEPs, it is likely that we missed many SEPs with antimicrobial activity – both SEPs and those encoded within larger genes. Finally, while our results suggest novel varieties of intra-and inter-species warfare, our ability to draw conclusions about the role SEPs might play in bacterial competition and microbial community assembly is limited. While we have experimental evidence of transcription and translation for five members of the list, for the majority of the 323 SEPs we do not have evidence of *in vivo* expression. Moreover, the activity that we do observe could be incidental and not related to biological function.

Taken together, our results have important implications for how we think about determinants of microbiome community structure and the ability of new invaders to establish a niche in a complex community. Our pipeline represents an important platform for the discovery of peptide antibiotics including those that might spare commensals. Here, we demonstrated the ability to find peptides that are as active as clinically relevant AMPs, such as polymyxin B. Future studies will help inform whether SEPs that are predicted to shape the microbiome are necessary and sufficient to do so in complex communities, and additional mining and drug development efforts are expected to identify SEPs that may have strong translational utility.

## Acknowledgments

Cesar de la Fuente-Nunez holds a Presidential Professorship at the University of Pennsylvania and acknowledges funding from the Procter & Gamble Company, United Therapeutics, a BBRF Young Investigator Grant, the Nemirovsky Prize, Penn Health-Tech Accelerator Award, and the Dean’s Innovation Fund from the Perelman School of Medicine at the University of Pennsylvania. Research reported in this publication was supported by the Langer Prize (AIChE Foundation), the National Institute of General Medical Sciences of the National Institutes of Health under award number R35GM138201, and the Defense Threat Reduction Agency (DTRA; HDTRA11810041, HDTRA1-21-1-0014, and HDTRA1-23-1-0001). This work was also supported by a Paul Allen Distinguished Investigator Award, NIH R01AI148623 and R01AI143757 (to A.S.B.). We thank Dr. Mark Goulian for kindly donating the following strains: *Escherichia coli* AIC221 [*Escherichia coli* MG1655 phnE_2::FRT (control strain for AIC222)] and *Escherichia coli* AIC222 [*Escherichia coli* MG1655 pmrA53 phnE_2::FRT (polymyxin resistant)]. We thank Dr. Andy Goodman for kindly donating wild type *Bacteroides thetaiotaomicron* ATCC 29741 (Background: VPI 5482). We thank the H-MARC core in the Center for Molecular Studies in Digestive and Liver Diseases (P30 DK050306) for kindly donating strains *Acinetobacter baumannii* ATCC 19606 and *Klebsiella pneumoniae* ATCC 13883. We thank the de la Fuente Lab members and Roby Bhattacharyya for insightful discussions. Figures created with BioRender.com are attributed as such. Molecules were rendered using the PyMOL Molecular Graphics System, Version 2.1 Schrödinger, LLC.

## Funding

National Institutes of Health grant R35GM138201 (CFN)

Defense Threat Reduction Agency grant HDTRA11810041 (CFN)

Defense Threat Reduction Agency grant HDTRA1-21-1-0014 (CFN)

Defense Threat Reduction Agency grant HDTRA1-23-1-0001 (CFN)

## Author contributions

Conceptualization: MDTT, EB, ASB, CFN

Methodology: MDTT, EB, AC, HS, CN

Investigation: MDTT, EB, AC

Visualization: MDTT, EB, AC, CN

Funding acquisition: ASB, CFN

Supervision: ASB, CFN

Software: EB, HS, CN

Formal analysis: MDTT, EB

Writing – original draft: MDTT, EB, ASB, CFN

Writing – review & editing: MDTT, EB, AC, CN, HS, ABS, CFN

## Competing interests

Cesar de la Fuente-Nunez provides consulting services to Invaio Sciences and is a member of the Scientific Advisory Boards of Nowture S.L. and Phare Bio. The de la Fuente Lab has received research funding or in-kind donations from United Therapeutics, Strata Manufacturing PJSC, and Procter & Gamble, none of which were used in support of this work. An invention disclosure associated with the work has been submitted.

## STAR Methods

### Key resource table

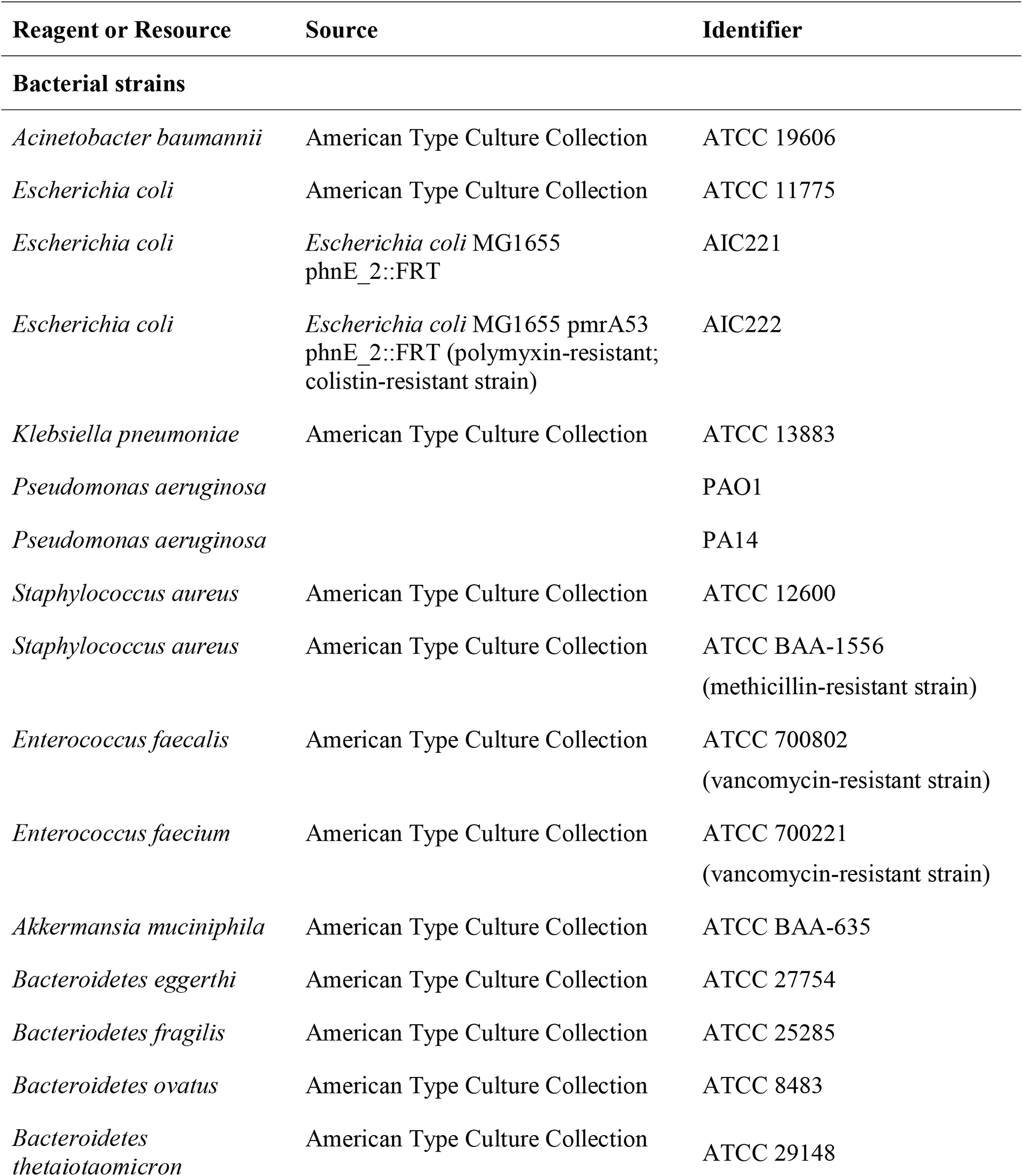
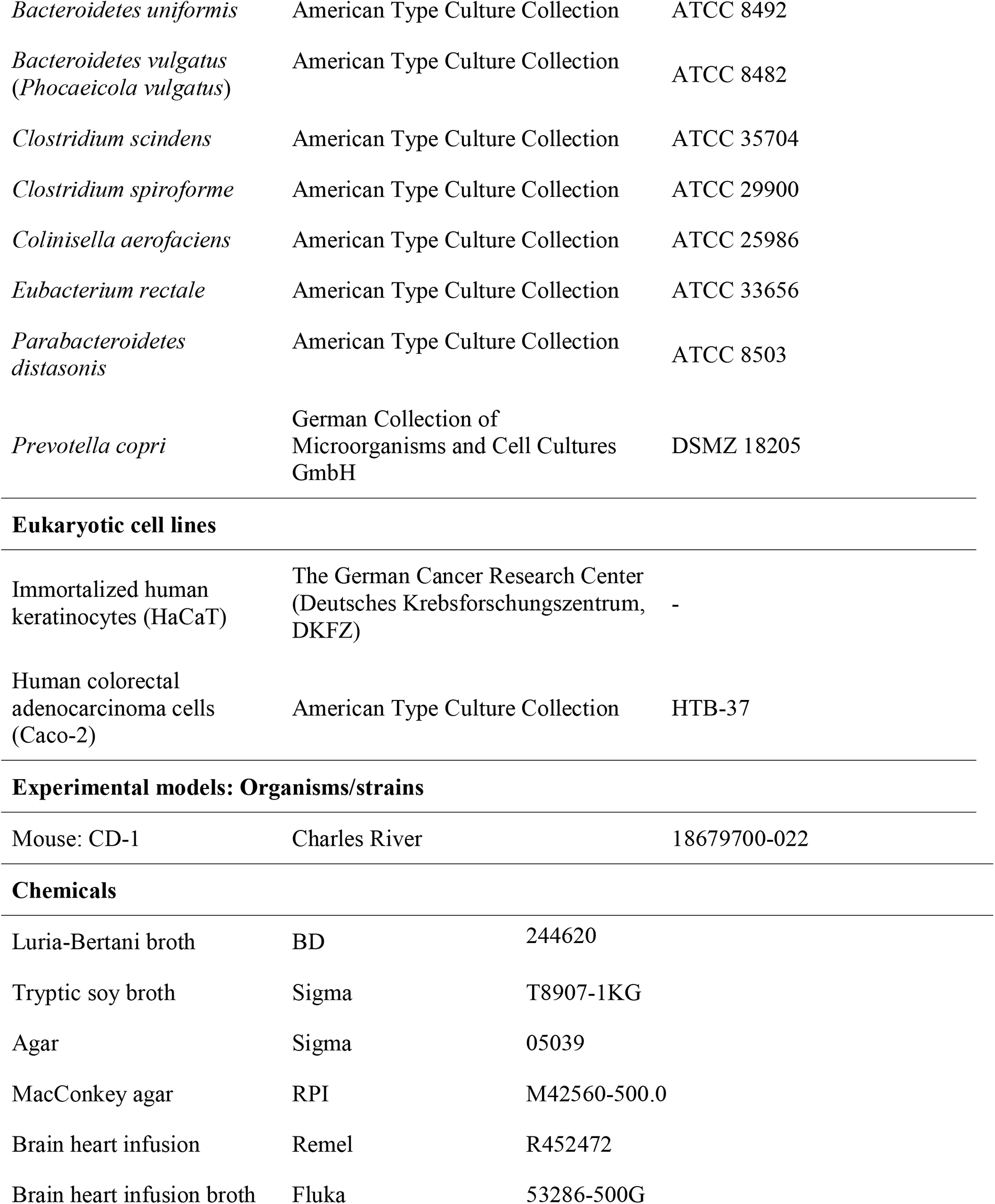
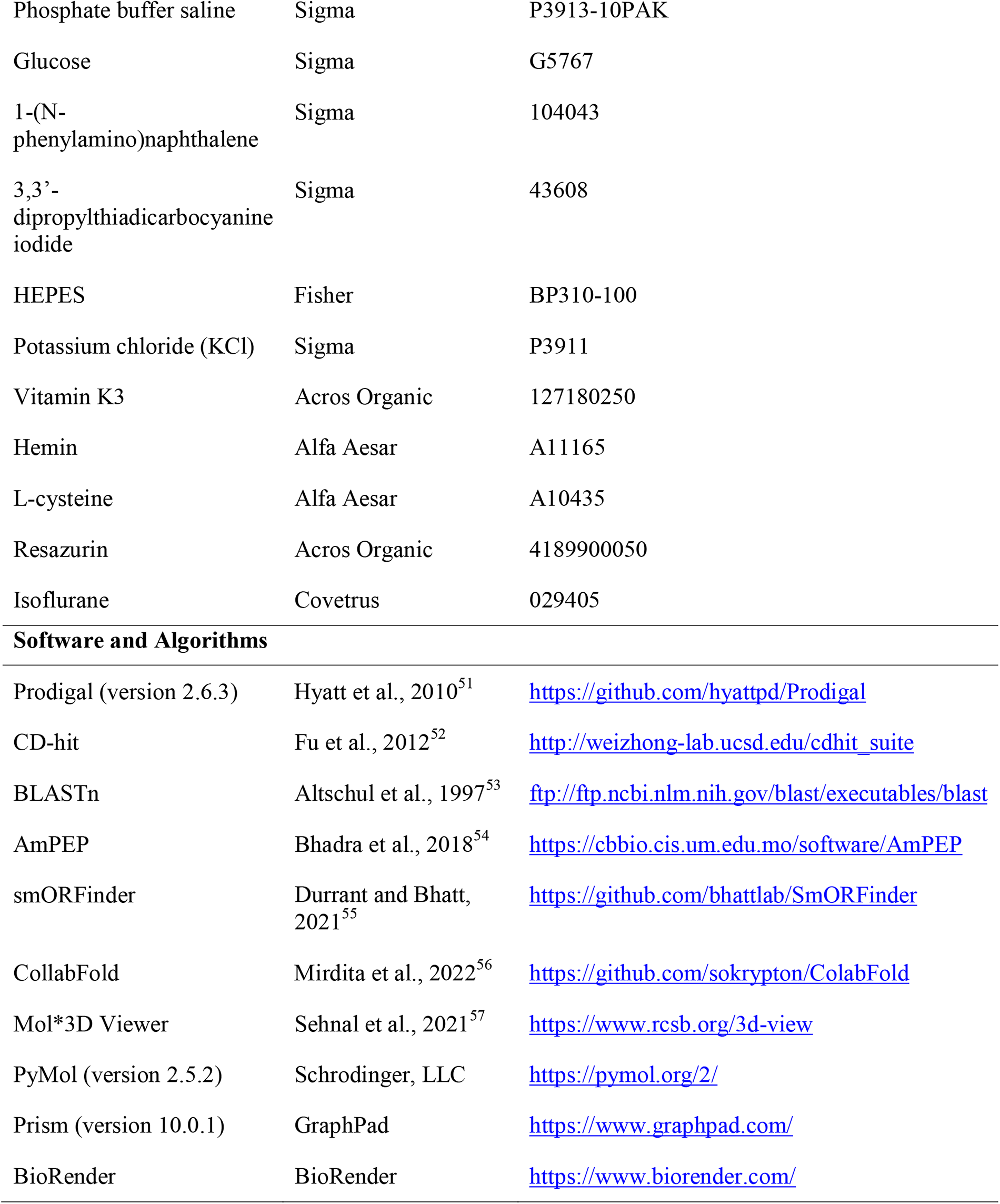

Key resources table
| Reagent or Resource | Source | Identifier |
| --- | --- | --- |
| **Bacterial strains** |  |  |
| *Acinetobacter baumannii* | American Type Culture Collection | ATCC 19606 |
| *Escherichia coli* | American Type Culture Collection | ATCC 11775 |
| *Escherichia coli* | *Escherichia coli* MG1655<br>phnE_2::FRT | AIC221 |
| *Escherichia coli* | *Escherichia coli* MG1655 pmrA53<br>phnE_2::FRT (polymyxin-resistant;<br>colistin-resistant strain) | AIC222 |
| *Klebsiella pneumoniae* | American Type Culture Collection | ATCC 13883 |
| *Pseudomonas aeruginosa* |  | PAO1 |
| *Pseudomonas aeruginosa* |  | PA14 |
| *Staphylococcus aureus* | American Type Culture Collection | ATCC 12600 |
| *Staphylococcus aureus* | American Type Culture Collection | ATCC BAA-1556<br>(methicillin-resistant strain) |
| *Enterococcus faecalis* | American Type Culture Collection | ATCC 700802<br>(vancomycin-resistant strain) |
| *Enterococcus faecium* | American Type Culture Collection | ATCC 700221<br>(vancomycin-resistant strain) |
| *Akkermansia muciniphila* | American Type Culture Collection | ATCC BAA-635 |
| *Bacteroidetes eggerthi* | American Type Culture Collection | ATCC 27754 |
| *Bacteroidetes fragilis* | American Type Culture Collection | ATCC 25285 |
| *Bacteroidetes ovatus* | American Type Culture Collection | ATCC 8483 |
| *Bacteroidetes*<br>*thetaiotaomicron* | American Type Culture Collection | ATCC 29148 |
| *Bacteroidetes uniformis* | American Type Culture Collection | ATCC 8492 |
| *Bacteroidetes vulgatus*<br>(*Phocaeicola vulgatus*) | American Type Culture Collection | ATCC 8482 |
| *Clostridium scindens* | American Type Culture Collection | ATCC 35704 |
| *Clostridium spiroforme* | American Type Culture Collection | ATCC 29900 |
| *Colinisella aerofaciens* | American Type Culture Collection | ATCC 25986 |
| *Eubacterium rectale* | American Type Culture Collection | ATCC 33656 |
| *Parabacteroidetes distasonis* | American Type Culture Collection | ATCC 8503 |
| *Prevotella copri* | German Collection of<br>Microorganisms and Cell Cultures<br>GmbH | DSMZ 18205 |
| **Eukaryotic cell lines** |  |  |
| Immortalized human<br>keratinocytes (HaCaT) | The German Cancer Research Center<br>(Deutsches Krebsforschungszentrum,<br>DKFZ) | - |
| Human colorectal<br>adenocarcinoma cells<br>(Caco-2) | American Type Culture Collection | HTB-37 |
| **Experimental models: Organisms/strains** |  |  |
| Mouse: CD-1 | Charles River | 18679700-022 |
| **Chemicals** |  |  |
| Luria-Bertani broth | BD | 244620 |
| Tryptic soy broth | Sigma | T8907-1KG |
| Agar | Sigma | 05039 |
| MacConkey agar | RPI | M42560-500.0 |
| Brain heart infusion | Remel | R452472 |
| Brain heart infusion broth | Fluka | 53286-500G |
| Phosphate buffer saline | Sigma | P3913-10PAK |
| Glucose | Sigma | G5767 |
| 1-(N-phenylamino)naphthalene | Sigma | 104043 |
| 3,3'-dipropylthiadicarbocyanine iodide | Sigma | 43608 |
| HEPES | Fisher | BP310-100 |
| Potassium chloride (KCl) | Sigma | P3911 |
| Vitamin K3 | Acros Organic | 127180250 |
| Hemin | Alfa Aesar | A11165 |
| L-cysteine | Alfa Aesar | A10435 |
| Resazurin | Acros Organic | 4189900050 |
| Isoflurane | Covetrus | 029405 |
| **Software and Algorithms** |  |  |
| Prodigal (version 2.6.3) | Hyatt et al., 2010<sup>51</sup> | https://github.com/hyattpd/Prodigal |
| CD-hit | Fu et al., 2012<sup>52</sup> | http://weizhong-lab.ucsd.edu/cdhit_suite |
| BLASTn | Altschul et al., 1997<sup>53</sup> | ftp://ftp.ncbi.nlm.nih.gov/blast/executables/blast |
| AmPEP | Bhadra et al., 2018<sup>54</sup> | https://cbbio.cis.um.edu.mo/software/AmPEP |
| smORFinder | Durrant and Bhatt, 2021<sup>55</sup> | https://github.com/bhattlab/SmORFinder |
| CollabFold | Mirdita et al., 2022<sup>56</sup> | https://github.com/sokrypton/ColabFold |
| Mol*3D Viewer | Sehnal et al., 2021<sup>57</sup> | https://www.rcsb.org/3d-view |
| PyMol (version 2.5.2) | Schrodinger, LLC | https://pymol.org/2/ |
| Prism (version 10.0.1) | GraphPad | https://www.graphpad.com/ |
| BioRender | BioRender | https://www.biorender.com/ |

### Resource Availability

#### Lead contact

Further information and requests for resources should be directed to and will be fulfilled upon reasonable request by the lead contact, Cesar de la Fuente-Nunez.

#### Materials availability

This study did not generate new unique reagents.

#### Data and code availability

Test results used for analysis will be provided upon reasonable request.

- HMPI-II metagenomes from which SEPs were annotated are publicly available from the NIH Human Microbiome Project: https://www.hmpdacc.org/hmasm2/.
- Any additional information required to reanalyze the data reported in this paper is available from the lead contact upon request.

### Experimental model

#### Bacterial strains and growth conditions

*Acinetobacter baumannii* ATCC 19606, *Escherichia coli* ATCC 11775, *Escherichia coli* AIC221 [*Escherichia coli* MG1655 phnE_2::FRT (control strain for AIC222)] and *Escherichia coli* AIC222 [*Escherichia coli* MG1655 pmrA53 phnE_2::FRT (polymyxin resistant; colistin-resistant strain)], *Klebsiella pneumoniae* ATCC 13883, *Pseudomonas aeruginosa* PAO1, *Pseudomonas aeruginosa* PA14, *Staphylococcus aureus* ATCC 12600, *Staphylococcus aureus* ATCC BAA-1556 (methicillin-resistant strain), *Enterococcus faecalis* ATCC 700802 (vancomycin-resistant strain), *Enterococcus faecium* ATCC 700221 (vancomycin-resistant strain) were grown and plated on Luria-Bertani (LB), Pseudomonas Isolation (*Pseudomonas aeruginosa* strains), and MacConkey (*A. baumannii* strain) agar plates and incubated overnight at 37 °C from frozen stocks. After incubation, one isolated colony was transferred to 5 mL of medium (LB), and cultures were incubated overnight (16 h) at 37 °C. The following day, inocula were prepared by diluting the overnight cultures 1:100 in 5 mL of the respective media and incubating them at 37 °C until bacteria reached logarithmic phase (OD_600_ = 0.3-0.5).

*Akkermansia muciniphila* ATCC BAA-635, *Bacteroidetes eggerthi* ATCC 27754, *Bacteriodetes fragilis* ATCC 25285, *Bacteroidetes ovatus* ATCC 8483, *Bacteroidetes thetaiotaomicron* ATCC 29148, *Bacteroidetes uniformis* ATCC 8492, *Bacteroidetes vulgatus* ATCC 8482 (*Phocaeicola vulgatus*), *Clostridium scindens* ATCC 35704, *Clostridium spiroforme* ATCC 29900, *Colinisella aerofaciens* ATCC 25986, *Eubacterium rectale* ATCC 33656, *Parabacteroidetes distasonis* ATCC 8503, and *Prevotella copri* DSMZ 18205 were the gut commensal strains used in this study. All the commensal microorganisms were cultured in brain heart infusion (BHI) broth and agar plates enriched with 0.1% (v:v) vitamin K3 (1 mg mL^-1^), 1% (v:v) hemin (1 mg mL^-1^, diluted with 10 mL of 1 N sodium hydroxide), and 10% (v:v) _L_-cysteine (0.05 mg mL^-1^), from cryopreserved stocks and incubated overnight at 37 °C. Resazurin was used as oxygen indicator. After the incubation period, a single isolated colony was transferred to 3 mL of BHI broth and incubated overnight at 37 °C. The next day, inocula were prepared by diluting the bacterial overnight cultures 1:100 in 3 mL of BHI broth and incubated at 37 °C until reaching the logarithmic phase (OD_600_ = 0.3-0.5).

#### Eukaryotic cell culture conditions

Immortalized human keratinocytes (HaCaT – obtained from DKFZ)^37^ and human colorectal adenocarcinoma cells (Caco-2 HTB-37, ATCC) were cultured in high-glucose Dulbecco’s modified Eagle’s medium (glumax DMEM – Gibco 11965092) supplemented with 1% v/v antibiotics (penicillin/streptomycin) and 10% v/v fetal bovine serum (FBS). HaCaT and Caco-2 cell lines were grown at 37 °C in a humidified atmosphere containing 5% CO_2_.

#### Skin abscess infection mouse model

To assess the effectiveness of the peptides against *A. baumannii* ATCC 19606, the bacteria were cultured in tryptic soy broth (TSB) medium until reaching an OD_600_ of 0.5. Subsequently, the cells were washed twice with sterile PBS (pH 7.4) and suspended to a final concentration of 5×10^6^ colony-forming units (CFU) mL^-1^. For the in vivo experiments, six-week-old female CD-1 mice were anesthetized with isoflurane and subjected to a superficial linear skin abrasion on their backs. A 20 μL aliquot containing the bacterial load suspended in PBS was then inoculated over the scratched area. The peptides, diluted in water at their MIC value, were administered to the infected area one hour after the infection. Two and four days post-infection, the animals were euthanized, and the skin area with the infection was excised and homogenized for 20 min using a bead beater (25 Hz). The homogenates were then 10-fold serially diluted for CFU quantification. The experiments were conducted with three mice per group (n=3), and a total of three independent experiments were performed using the MIC values of the peptides (n=9). The skin abscess infection mouse model was revised and approved by the University Laboratory Animal Resources (ULAR) from the University of Pennsylvania (Protocol 806763).

#### Deep thigh infection mouse model

To induce neutropenia in mice, two intraperitoneal doses of cyclophosphamide (150 mg Kg^-1^) were administered at a 72-hour interval. One day after the last cyclophosphamide dose, the mice were intramuscularly infected in their right thigh with a bacterial load of 5×10^6^ CFU mL^-1^ of *A. baumannii* ATCC 19606, which had been cultured in tryptic soy broth (TSB), washed twice with PBS (pH 7.4), and resuspended to the required concentration. Two hours after infection, the mice were treated with peptides suspended in water via intraperitoneal injection. Before each injection, the mice were anesthetized with isoflurane and their respiratory rate and pedal reflexes were monitored. Subsequently, we closely monitored the progression of the infection and euthanized the mice accordingly. The infected area was removed two and four days post-infection, homogenized for 20 min (25 Hz) using a bead beater, and 10-fold serially diluted for CFU quantification on MacConkey agar plates. Each experimental group consisted of 3 mice (n=3), and a total of three independent experiments were conducted, resulting in a sample size of 9 mice per condition (n=9). However, during one of the replicates, an unexpected bacterial growth occurred suddenly, leading to a significant increase in bacterial counts by three orders of magnitude (inoculum 5×10^6^ CFU mL-1 and counts ∼10^9^ CFU mL-1) within a single day. As a consequence, all three mice in that replicate perished (n=3), which accounts for the variation in the number of mice (n=6) observed on day 8 for each condition. The deep thigh infection mouse model was revised and approved by the University Laboratory Animal Resources (ULAR) from the University of Pennsylvania (Protocol 807055).

### Methods details

#### Generation of list of 444,054 smORF family representatives from multiple human associated metagenomes

We started with the list of 444,054 family representatives previously reported in Sberro et al.^18^. The computational methodology is summarized here. Contigs from 1,773 HMPI-II human-associated metagenomes from four major body sites were downloaded. All ORFs ≥15 bp were predicted using MetaProdigal^51^. The list was then filtered to only contain small ORFs ≤150 bp, resulting in a set of 2,514,099 smORFs. The proteins encoded by these smORFs were clustered into families using the following parameters: -n 2 -p 1 -c 0.5 -d 200 -M 50000 -l 5 -s 0.95 –aL 0.95 –g 1 (family members required to have 40-50% homology; shorter sequences required to be ≥ 95% length of the cluster representative; and the alignment was required to cover ≥ 95% of the longer sequence). This resulted in 444,054 clusters, each of which was assigned a ‘cluster representative’ by CD-Hit^52^. The cluster representative for each of the 444,054 clusters was used in subsequent parts of our analysis. Hereinafter, we use the family ID interchangeably to refer to the family representative.

#### Antimicrobial peptide prediction

AmPEP^58^ was applied (default parameters) on the 444,054 representatives of families. We considered all peptides with AmPEP score greater than 0.5 generating a list of 11,710 family representatives with likelihood of antimicrobial activity.

#### Application of smORFinder to predict high confidence peptides

SmORFinder^59^ was run on all metagenomes from the Human Microbiome Project (HMPI-II)^60^ as previously reported. We filtered the list of 444,054 family representatives for those that were also called by smORFinder. This resulted in a list of 38,965 family representatives. We then took the intersection of the list of 38,965 representatives called by smORFinder and the list of 11,710 antimicrobial representatives called by AmPEP. This yielded the list of 323 representatives that were both smORFinder positive and had AmPEP score ≥0.5. This list was used in our subsequent analysis.

#### Inclusion and exclusion criteria to select peptides for activity testing

Our approach to selecting peptide sequences from the list of 323 was as follows. First, we applied the following exclusion criteria: (1) we removed from consideration all peptides with more than one cysteine residues, these were considered undesirable candidates due to the tendency of these peptides to oxidize and cross-link aggregating in solution; (2) we removed peptides that have a high mean hydrophobicity owing to the difficulty of chemical synthesis and the tendency of these peptides to aggregate. (3) We chose peptides representing the “known” sequence space, these were peptides that shared many features of known AMPs and looked to have high confidence of being antimicrobial. (4) We chose peptides representing the negative sequence space, these were peptides that shared similarity to sequences that are either known to not be antimicrobial or not likely to be antimicrobial (*e.g.*, net negative charge). (5) We chose peptides that represented the edge of the “known” sequence space. These were peptides that looked promising (*e.g.*, appear to form α-helices, or contain positive charge or poly-lysine residues), but contained some novelty. (6) We chose peptides that represented the “unexplored” sequence space that would typically be looked over by conventional structure-activity relationship design approaches. These were peptides with unconventional sequences, for example ones that have amino acid residues that are not typically found in AMPs, such as polar uncharged residues (Asn, Gln, Met, Ser, Thr).

In all cases, we chose to test only the representative member of the family. For promising candidates, there is the option ’expand’ the family to test the other sequences in the cluster. Given that the families were clustered at the 40-50% homology threshold, we expect peptides within the same family to have dramatic differences in activity. In these cases, we could look at individual members of a family for conserved sites of activity. In one case (family 420019), where the cluster representative could not be synthesized, we took the consensus sequence of the alignment of all 47 family members in family 420019.

#### Peptide synthesis

All peptides used in the experiments were purchased from AAPPTec and synthesized by solid-phase peptide synthesis using the Fmoc strategy.

#### Minimal inhibitory concentration determination

The 78 SEPs underwent broth microdilution assays to evaluate their *in vitro* antimicrobial activity. The determination of the minimum inhibitory concentration (MIC) values was done by utilizing the broth microdilution technique, wherein a starting inoculum of 2×10^6^ cells was introduced into LB, in nontreated polystyrene microtiter 96-well plates. Peptides, prepared as aqueous solutions, were added to the plate and two-fold diluted (1 to 128 µmol L^-1^). The MIC was identified as the lowest concentration of peptide that completely inhibited the visible growth of bacteria after 20 h of incubation at 37 °C. The plates were then analysed using a spectrophotometer at 600 nm. The assays were conducted in triplicate to ensure statistical reliability.

#### Circular dichroism assays

Circular dichoism assays were performed as previously described (CITE). Briefly, the experiments were conducted at the University of Pennsylvania’s Biological Chemistry Resource CEnter (BCRC) using a J1500 circular dichroism spectropolarimeter (Jasco). The experiments were carried out at 25 °C, and the circular dichroism spectra represent an average of three accumulations. Three spectra accumulations were obtained using a quartz cuvette with an optical path length of 1.0 mm, covering a wavelength range from 260 to 190 nm at a rate of 50 nm min^-1^ and a bandwidth of 0.5 nm. The concentration of all peptides tested was 50 μmol L^-1^, and the measurements were performed in a mixture of water and trifluoroethanol (TFE) in a 3:2 ratio. Respective baselines were recorded before taking the measurements, and a Fourier transform filter was applied to minimize background effects. Helical fraction values were calculated using the single spectra analysis tool on the server BeStSel^29^.

#### Synergy assays

The selection of *A. baumannii* ATCC 19606 for the synergy assays was based on its significance as a pathogen that exhibits intrinsic resistance to antimicrobial agents and its capacity to infect various sites including the urinary tract, gastrointestinal tissue, and skin and soft tissues^43^. Following determination of the minimum inhibitory concentration (MIC) for each peptide, the most efficacious SEPs against *A. baumannii* were subjected to orthogonal dilution and concentration range from 2×MIC to 0.03×MIC. The plates were then incubated at 37 °C for 24 h. All assays were conducted in triplicate to ensure reliability of results.

#### Cytoplasmic membrane depolarization assays

The determination of the peptides’ ability to depolarize the cytoplasmic membrane was carried out by measuring the fluorescence of the membrane potential-sensitive dye, 3,3’-dipropylthiadicarbocyanine iodide (DiSC_3_-5). For this experimental protocol, we cultured *A. baumannii* ATCC 19606, *E. coli* ATCC 11775, *E. coli* AIC221, *E. coli* AIC222, *K. pneumoniae* ATCC 13883, *P. aeruginosa* PA14, *S. aureus* ATCC 12600, methicillin-resistant *S. aureus* ATCC BAA-1556, vancomycin-resistant *E. faecalis* ATCC 700802, and vancomycin-resistant *E. faecium* ATCC 700221 with agitation at 37 °C until OD_600_ = 0.5. Subsequently, the cells were centrifuged and washed twice with washing buffer (20 mmol L^-1^ glucose, 5 mmol L^-1^ HEPES, pH 7.2) and diluted 10-fold in the same buffer containing 0.1 mol L^-1^ KCl. Next, the cells (100 µL) were incubated for 15 min with DiSC_3_-5 (20 nmol L^-1^) until the fluorescence stabilization. We then incubated the peptides (100 µL solution at MIC values) and the changes in fluorescence emission intensity of DiSC_3_-5 (λ_ex_ = 622 nm, λ_em_ = 670 nm) were used to track depolarization of the cytoplasmic membrane for 60 min. The relative fluorescence was calculated using a non-linear fit and the positive control (buffer + bacteria + fluorescent dye + polymyxin B) served as baseline for comparison. The following equation was applied to determines the % difference between the baseline and the sample:

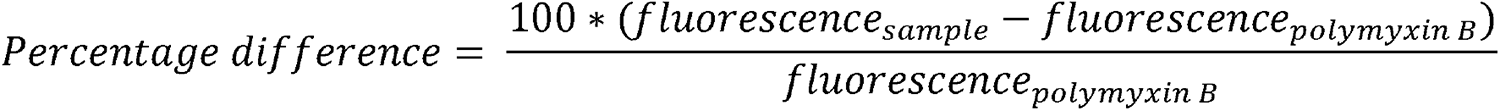

#### Outer membrane permeabilization assays

The membrane permeability of SEPs was assessed using the 1-(N-phenylamino)naphthalene (NPN) uptake assay. *A. baumannii* ATCC 19606, *E. coli* ATCC 11775, *E. coli* AIC221, *E. coli* AIC222, *K. pneumoniae* ATCC 13883, and *P. aeruginosa* PA14 were cultured until reaching an OD_600_ of 0.4. The cells were then centrifuged (10,000 rpm at 4 °C for 10 min), washed, and resuspended in a buffer (5 mmol L^-1^ HEPES, 5 mmol L^-1^ glucose, pH 7.4). Four µL of NPN solution (0.5 mmol L^-1^) was added to the bacterial solution in a white 96-well plate (total volume of 100 µL). The baseline fluorescence, i.e., fluorescence before the addition of peptides, was measured at λ_ex_ = 350 nm and λ_em_ = 420 nm. Membrane permeability was monitored after the addition of peptides at their MIC (100 µL) until no further increase was observed (for 45 min). The relative fluorescence was calculated using a non-linear fit and the positive control (buffer + bacteria + fluorescent dye + polymyxin B) was used as the baseline. The following equation was used to determine the % difference between the baseline and the sample:

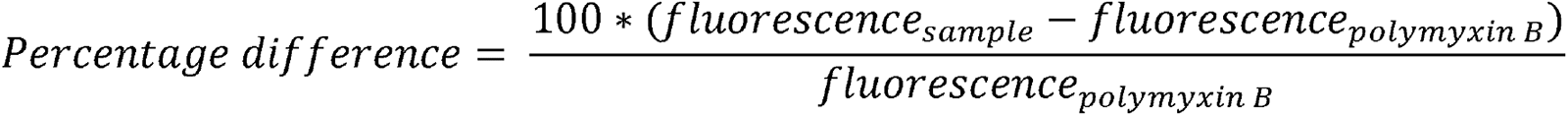

#### Cytotoxicity assays

Cells were seeded into 96-well plates at the density of 5×10^3^ cells per well for 24 h. After this period, we treated them with increasing concentrations of peptide (4-64 μmol L^-1^). Next, MTT reagent at 0.5 mg mL^-1^ in DMEM medium without phenol red was used to replace cell culture supernatant (100 μL per well). The samples were incubated for 4 h at 37 °C to obtain the insoluble formazan salts. Then, the resulting salts were solubilized in 0.04 mol L^-1^ HCl in anhydrous isopropanol and quantified using a spectrophotometer at 570 nm.

### Quantification and statistical analysis

#### Reproducibility of the experimental assays

All assays were performed in three independent biological replicates. Cytotoxic activity values were determined through non-linear regression analysis, utilizing a peptide gradient of concentrations. These values represent the concentrations required to lyse and kill 50% of the cells in the experiment. For the cytotoxic activity assays, two technical replicates were conducted within each of the three biological replicates. In the skin abscess and deep thigh infection mouse models, we employed three mice per group in three independent replicates, adhering to established protocols approved by the University Laboratory of Animal Resources (ULAR) at the University of Pennsylvania.

#### Statistical tests

In the mouse experiments, the statistical significance was determined using one-way ANOVA followed by Dunnett’s test. All the p values are shown for each of the groups, all groups were compared to the untreated control group.

#### Statistical analysis

All calculation and statistical analyses of the experimental data were conducted using GraphPad Prism v.10.0.1. Statistical significance between different groups was calculated using the tests indicated in each figure legend. No statistical methods were used to predetermine sample size.

