## Supplementary material for "Human gut metagenomic mining reveals an untapped source of peptide antibiotics": Torres_2023_SI.docx

**This PDF file includes:**

### Figures S1 to S6

### Table S1

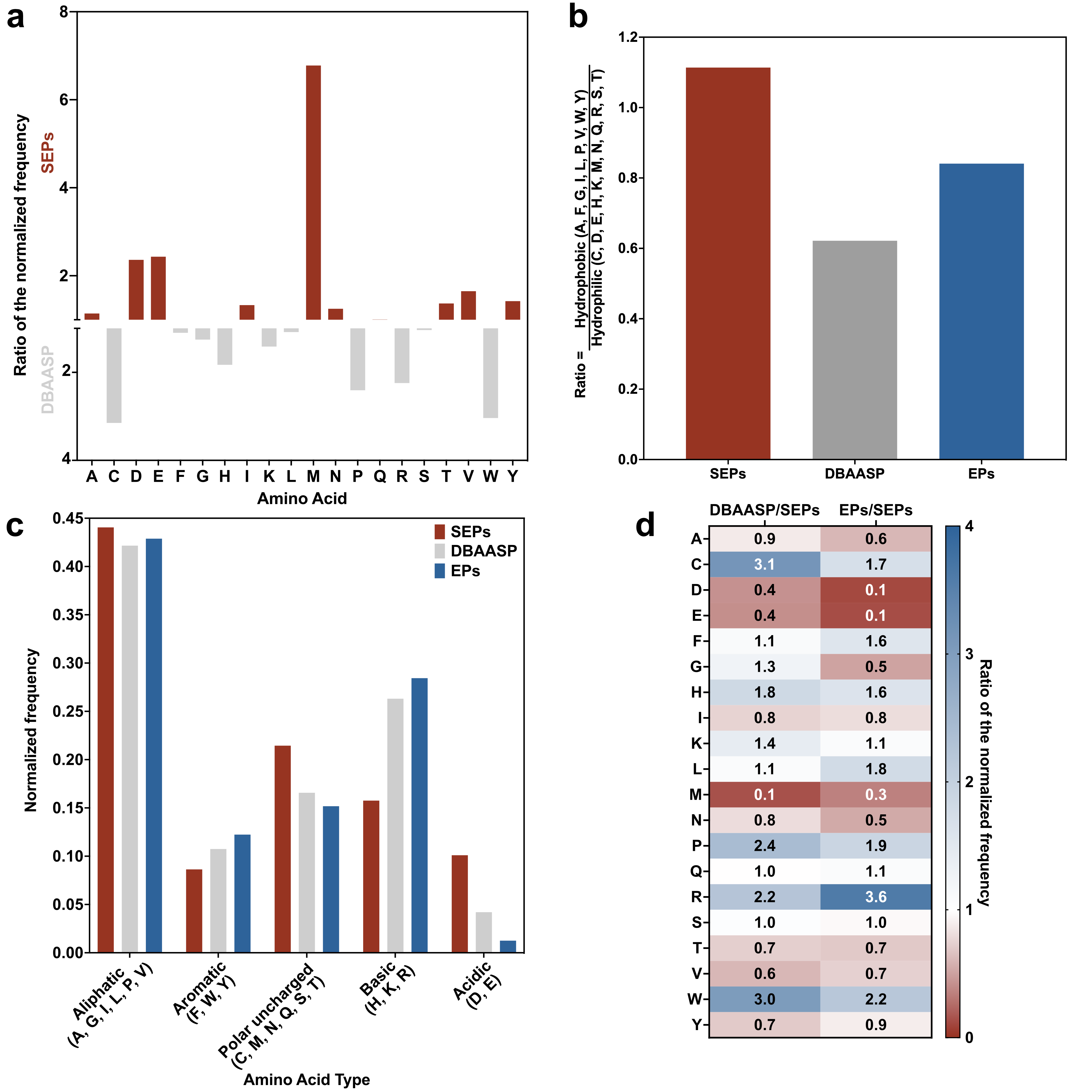

### Figure S1. Frequency distribution by amino acid and amino acid type of SEPs compared to known AMPs and EPs from the human proteome. a) Ratio of the normalized frequency of SEPs compared to AMPs from DBAASP. b) Ratio between Hydrophilic and Hydrophobic amino acid residues for each one of the three different classes of peptides (SEPs, AMPs, and EPs). SEPs are more hydrophobic than the other two classes. c) Normalized frequency of amino acid type showing that the SEPs present more negatively carged and polar uncharged residues than known AMPs and EPs, whereas AMPs and EPs have more positively charged and aromatic residues than EPs. d) Heat map showing the exact ratio of normalized frequency for each amino acid residue in comparisons between DBAASP and SEPs and EPs and SEPs. All 323 SEP and 43,000 EP candidates were used for the frequency calculation.

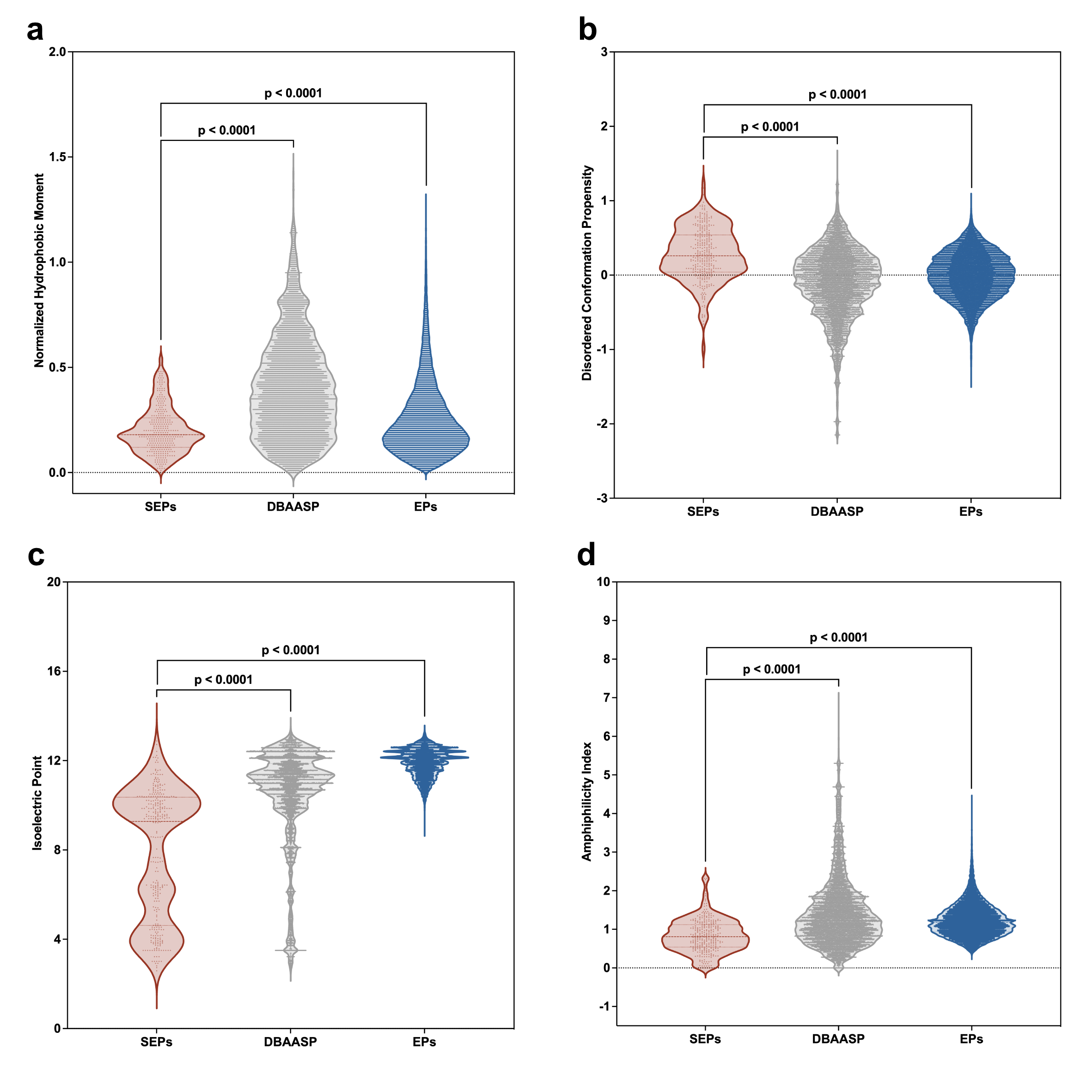

**Figure S2.** **Physicochemical features of smORF-encoded peptides (SEPs) compared to antimicrobial peptides and encrypted peptides from the human proteome.** The following physicochemical features were estimated using the Database of Antimicrobial Activity and Structure of Peptides (DBAASP) server^1^: **(a)** normalized hydrophobic moment, **(b)** disordered conformation propensity, **(c)** isoeletric point, and **(d)** amphiphilicity index. Those physicochemical features summed to net charge (**Figure 2b**) and normalized hydrophobic moment (**Figure 2c**) are the most relevant easy to extract properties that influence the antimicrobial activity and toxicity of peptides with antimicrobial properties.

**Table S1. Sequence and physicochemical features of the active SEPs.** Physicochemical features of the SEPs were calculated using the Database of Antimicrobial Activity and Structure of Peptides (DBAASP) server^1^.

| **SEP** | **Sequences** | **Lenght** | **Net Charge** | **Normalized Hydrophobicity** | **Normalized Hydrophobic Moment** | **Disordered Conformation Propensity** | **Isoelectric Point** | **Amphiphilicity Index** |
| --- | --- | --- | --- | --- | --- | --- | --- | --- |
| Staphylococcin-1 | MQKLAEAIANTVKAGQDHDWAKLGTSIVGIAENGISLLGKVFGF | 44 | 0 | -0.23 | 0.18 | 0.32 | 7.47 | 0.64 |
| Caudoviralesin-1 | MSVQSRIRKIEKKKGMYKRRKEDIIKCKLKKQEKEN | 36 | 10 | 0.65 | 0.24 | -0.55 | 10.9 | 1.74 |
| Fusobacticin-1 | MKIVVKRVPKFLRGFVKLIFGLKDEK | 26 | 6 | 0.01 | 0.47 | 0.08 | 11.25 | 1.08 |
| Subdogranulin-1 | MLIKNIKWDTDGDMEALASLPIKRVP | 26 | 0 | -0.04 | 0.1 | 0.2 | 6.39 | 0.83 |
| Dysosmycin-1 | MTMNKFMADVDDLIFDGQEQSHSER | 25 | -4 | 0.14 | 0.19 | 0.15 | 3.86 | 0.5 |
| Veillonellin-1 | MVAAIVKAILAALTGLANALLPLFK | 25 | 2 | -0.57 | 0.4 | 0.7 | 10.69 | 0.29 |
| Bacteroidin-1 | MKTMKNFIQKVFKGDPMIWGYVMLS | 25 | 3 | -0.17 | 0.35 | 0.15 | 10.39 | 1.12 |
| Selenomin-1 | MADIFLLLLIFLMVYYVFGDELFPR | 25 | -2 | -0.56 | 0.18 | 0.78 | 3.57 | 0.55 |
| Staphylococcin-2 | MNLGKGISEAYKGFMKGAFGKNWNK | 25 | 4 | 0 | 0.2 | -0.13 | 10.59 | 1.26 |
| Prevotellin-1 | MSIAIVAIMIAQAVNLGNKINLGNK | 25 | 2 | -0.35 | 0.14 | 0.44 | 10.69 | 0.34 |
| Clostridisin-5 | MANGIIIIDKPAGWTKLSKKGKALF | 25 | 4 | -0.17 | 0.09 | 0.13 | 10.85 | 1.01 |
| Gemellin-1 | MSIYAHVTEKQRDNMADKFAKFMAL | 25 | 1 | 0.05 | 0.17 | 0.09 | 9.4 | 0.9 |
| Ruminococcin-1 | MVKLILLALFVWVIWLTLRKIFRGY | 25 | 4 | -0.47 | 0.2 | 0.54 | 11.42 | 1.25 |
| Fusobacticin-2 | MNYRLLVEIKEDKFIIKALSIGHRR | 25 | 3 | 0.11 | 0.18 | 0.06 | 10.5 | 1.1 |
| Faecalibacticin-1 | MKARIPKHREFIINFPDSIDQNKAN | 25 | 2 | 0.23 | 0.19 | -0.13 | 10.38 | 0.8 |
| Clostridisin-6 | MKKAIYRILILMVVAVMVVLSVLHF | 25 | 3 | -0.53 | 0.27 | 0.72 | 10.79 | 0.65 |
| Acholeplasin-1 | MKGIDASYEYEGYYTYKMEDLDKM | 24 | -3 | 0.1 | 0.22 | 0.04 | 3.93 | 1.67 |
| Rothiasin-1 | MKKLIVLALVGAAAFFVYKKKFAA | 24 | 5 | -0.37 | 0.15 | 0.4 | 10.9 | 0.98 |
| Keratinobacin-1 | MTIVVKKVPKFLRGFVKLIFGIKD | 24 | 5 | -0.2 | 0.54 | 0.26 | 11.22 | 0.87 |
| Streptocin-1 | MAKKKYQTPKVSELKSAAGINVKA | 24 | 5 | 0.13 | 0.04 | -0.14 | 10.73 | 1.23 |
| Nisserin-1 | MTYGLLLIAAIGALFAYKMHQQAK | 24 | 2 | -0.37 | 0.18 | 0.39 | 9.93 | 0.89 |
| Caudoviralesin-2 | MQTQKARKRELSVEEMIKRMQFDV | 24 | 2 | 0.4 | 0.24 | -0.15 | 10.36 | 1.08 |
| Fusobacticin-2 | MLPLISIVVGQVVANQITNLIKKD | 24 | 1 | -0.31 | 0.2 | 0.47 | 9.69 | 0.41 |
| Flavonin-1 | MASAAVVYITGGLTVPCVLRWMTR | 24 | 2 | -0.33 | 0.25 | 0.47 | 9.61 | 0.7 |
| Unknowing-1 | MAVKVAINGFGRIGRLAFRQMFGA | 24 | 4 | -0.19 | 0.48 | 0.26 | 12.41 | 0.51 |
| Prevotellin-2 | MLNYLYDRDINRYRAIIKALGLRK | 24 | 4 | 0.21 | 0.21 | -0.05 | 10.5 | 1.35 |
| Alistipin-1 | MKIGLVDVDGHHFPNLALMKLSA | 23 | 0 | -0.26 | 0.06 | 0.38 | 7.66 | 0.45 |
| Siphovirin-1 | MRITFKPLWKLLIDRDMTREDVR | 23 | 2 | 0.27 | 0.16 | -0.01 | 10.59 | 1.1 |
| Ehrlicin-1 | MEERENRIEIDEILKNKDKNKDK | 23 | -1 | 0.67 | 0.18 | -0.36 | 4.86 | 1.29 |
| Rothiasin-2 | MKKILCIAILAGAAYFGYKKSVA | 23 | 4 | -0.34 | 0.17 | 0.36 | 10.19 | 1.08 |
| Bacteroidin-2 | MGNLVAIVGRPNVGKSTLFNRFH | 23 | 3 | -0.1 | 0.27 | 0.15 | 12.12 | 0.44 |
| Cardiobacin-1 | MAKTYYDILGVAQNASAADIKKA | 23 | 1 | -0.09 | 0.28 | 0.19 | 9.28 | 0.97 |
| Faecalibacticin-2 | MQILTHLIFLLNLLKTLINHFS | 22 | 1 | -0.44 | 0.34 | 0.54 | 9.86 | 0.36 |
| Eubacticin-1 | MILKYREGGCTVGSGRVATVIE | 22 | 1 | -0.11 | 0.09 | 0.29 | 8.56 | 0.73 |
| Faecalibacticin-3 | MQIVVVKCPHALRWLLRAVFKV | 22 | 4 | -0.22 | 0.46 | 0.36 | 11.39 | 0.99 |
| Fastidin-1 | MEGYGNDNPGAKSQGIKQFMAK | 22 | 1 | 0.1 | 0.15 | -0.13 | 9.4 | 0.9 |
| Kingellin-1 | MTTPKFDRLAITQAVLLKYTYF | 22 | 2 | -0.14 | 0.11 | 0.22 | 9.93 | 0.96 |
| Ruminococcin-2 | MTRLKKVLAALIAVVFAALGLN | 22 | 3 | -0.41 | 0.23 | 0.57 | 11.56 | 0.44 |
| Ruminococcin-3 | MGIMGVIVLIILVIGVTKYGIR | 22 | 2 | -0.64 | 0.21 | 0.76 | 10.4 | 0.51 |
| Faecalibacticin-4 | MEMQYFKEYSPALGREMECKVY | 22 | -1 | 0.09 | 0.23 | 0.07 | 4.62 | 1.42 |
| Roseburin-1 | MAKTKVIAVANQKGGVGKSTTV | 22 | 4 | -0.08 | 0.03 | 0.09 | 11.15 | 0.72 |
| Lachnoanin-1 | MSYKQENFEEWIILIPVKIEY | 21 | -2 | -0.15 | 0.16 | 0.28 | 4.16 | 1.46 |
| Intestinin-1 | MAGGVVLIGITLVILFSSLFG | 21 | 0 | -0.8 | 0.13 | 0.93 | 3.5 | 0 |
| Clostridisin-7 | MTMRLLIAEDEADLAEARTVF | 21 | -3 | -0.07 | 0.17 | 0.47 | 3.8 | 0.41 |
| Myoviridin-1 | MGHINPWVAWMIEMLMFLHG | 20 | -1 | -0.54 | 0.17 | 0.53 | 6.05 | 0.9 |
| Aggregatin-1 | MLDVIIRIAEAVIDELKKNK | 20 | 0 | -0.03 | 0.51 | 0.33 | 6.49 | 0.8 |
| Siphovirin-2 | MCRVIAVSNQKGGVGKTVSC | 20 | 3 | -0.07 | 0.14 | 0.2 | 9.74 | 0.55 |
| Subdogranulin-1 | MPGQVEITVIRETRAVSYAK | 20 | 1 | 0.02 | 0.2 | 0.16 | 9.5 | 0.87 |
| Dysosmobin-1 | MGKKKTPQAKAYGAALQKG | 19 | 5 | 0.17 | 0.24 | -0.32 | 10.9 | 1.36 |
| Leptotricin-1 | MGILKKLWDMLPEGRPV | 17 | 1 | -0.1 | 0.34 | 0.18 | 9.69 | 1.06 |
| Myoviridin-2 | MLGAILGDIVGSPYEYG | 17 | -2 | -0.47 | 0.25 | 0.56 | 2.92 | 0.67 |
| Bacteriophagin-1 | MPEIEGIYYETEEDYY | 16 | -6 | -0.02 | 0.19 | 0.27 | 2.73 | 1.66 |
| Lachnoanin-2 | MVGATLIPLLSGLFG | 15 | 0 | -0.7 | 0.19 | 0.78 | 3.5 | 0 |

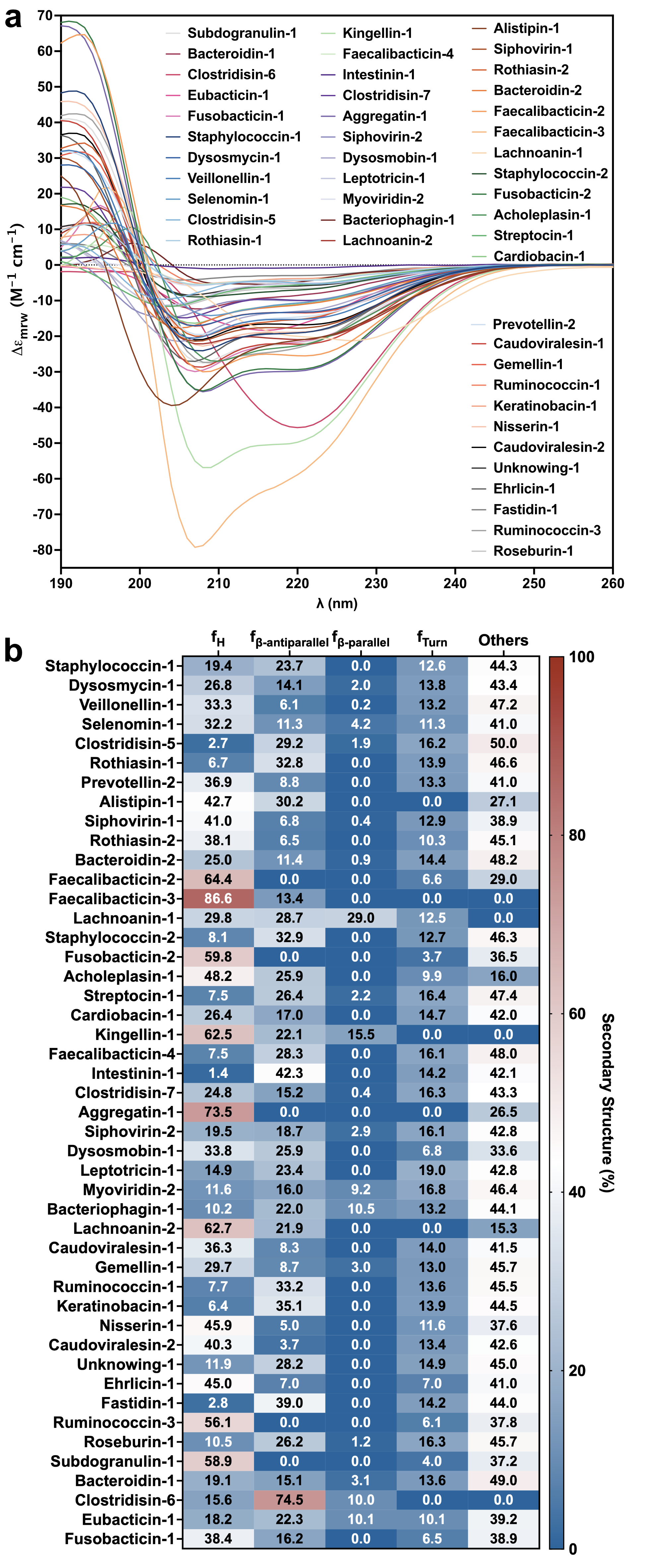

### Figure S3. Experimental determination of the secondary structure of SEPs. a) Circular Dichroism spectra of all active SEPs in TFE/water (3:2, v:v) and b) their secondary structure fractions calculated using the BeStSel server.

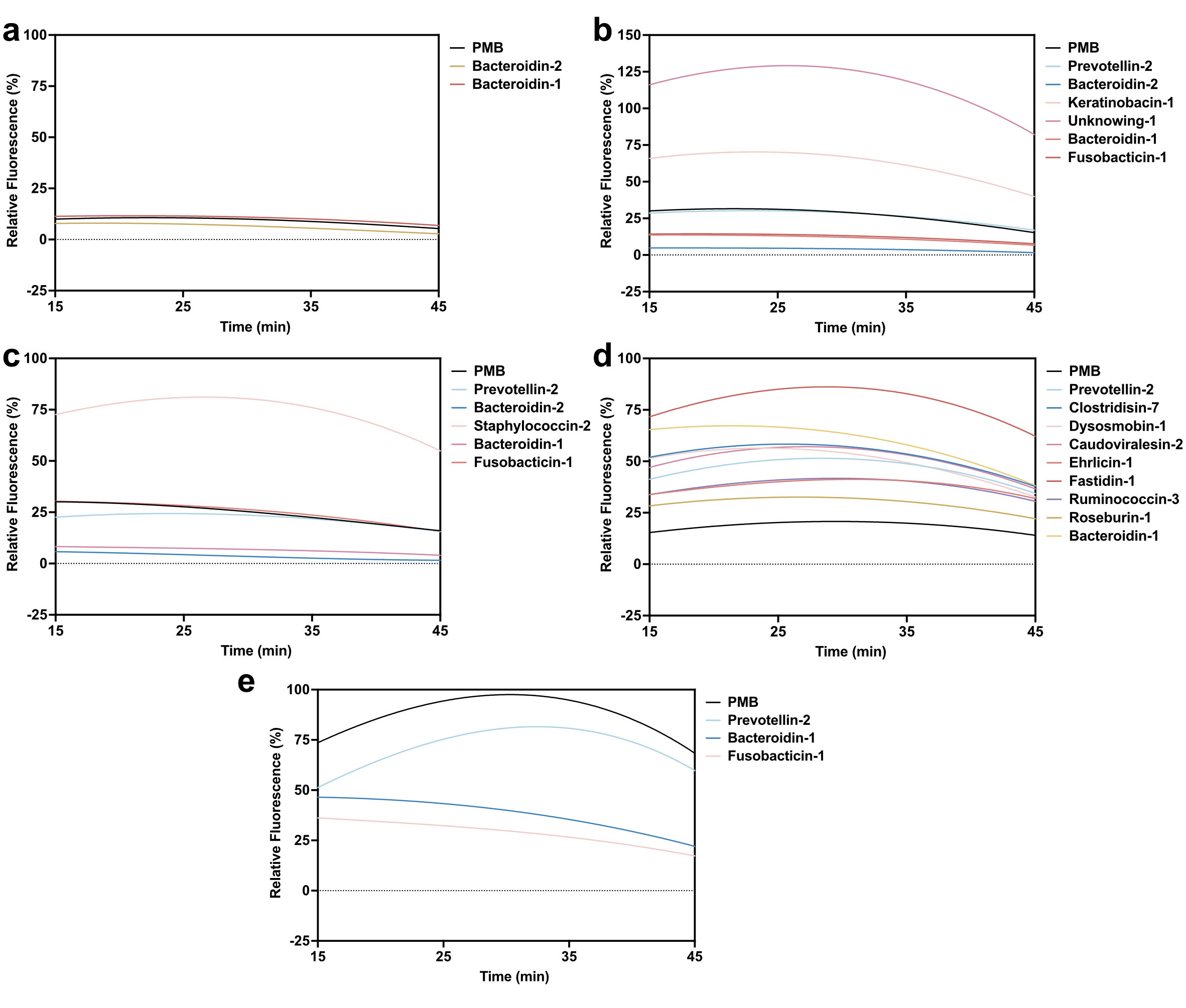

**Figure S4.** **Outer membrane permeabilization caused by SEPs on the membranes of Gram-negative pathogenic strains.** Permeabilization of the outer membrane using the probe 1-(N-phenylamino)naphthalene (NPN) on all the Gram-negative strains that were targeted by the SEPs: **(a)** *E. coli* ATCC 11775, **(b)** *E. coli* AIC221, **(c)** *E. coli* AIC222, **(d)** *K. pneumoniae* ATCC 13883, and **(e)** *P. aeruginosa* PA14. Polymyxin B was used as positive control and buffer with NPN and bacteria were used as baseline for the calculation of the relative fluorescence values.

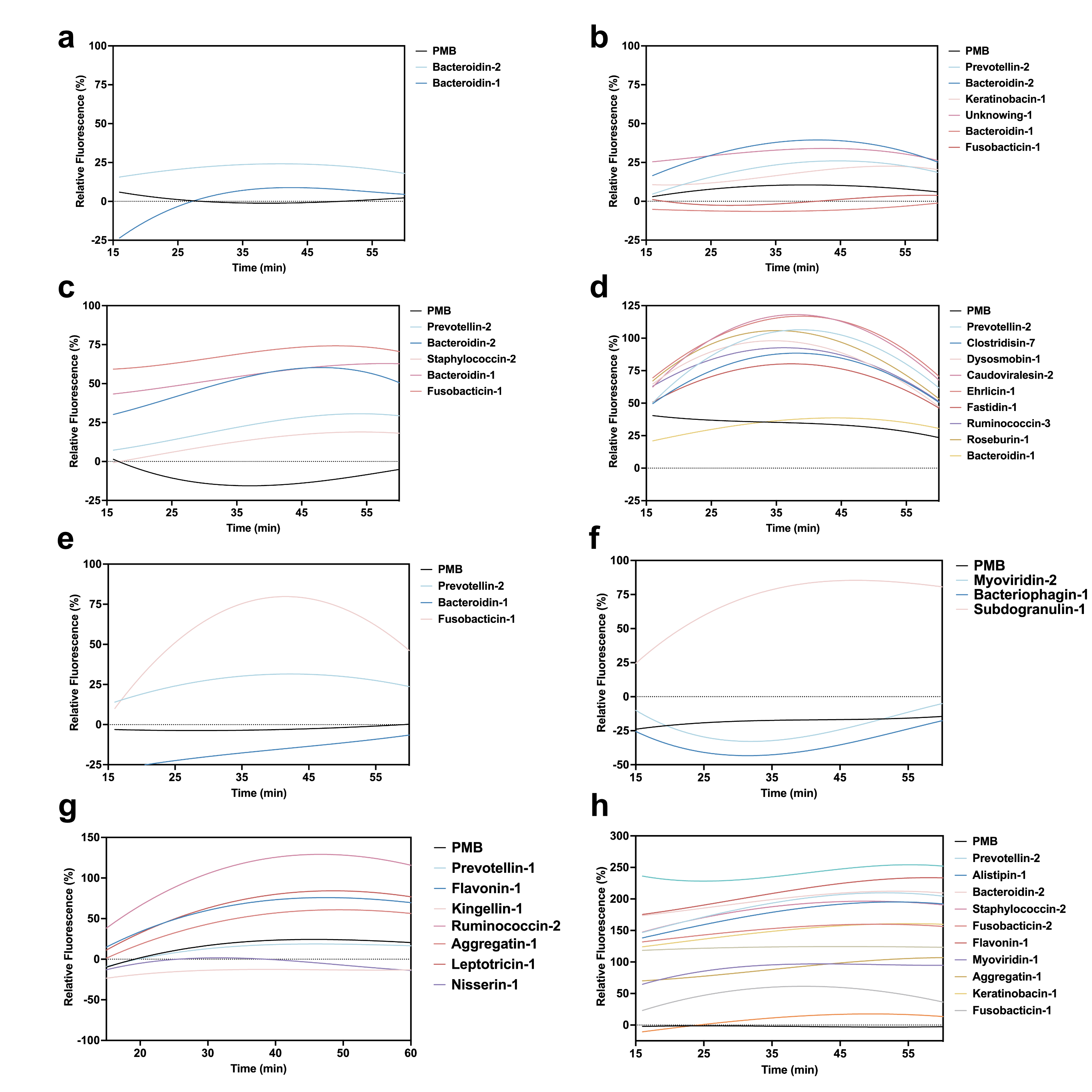

**Figure S5.** C**ytoplasmic membrane depolarization triggered by encrypted peptides from extinct organisms on *A. baumannii* and *P. aeruginosa* cell membranes.** Depolarization assays with the hydrophobic probe 3,3′-dipropylthiadicarbocyanine iodide (DiSC_3_-5) on all pathogenic strains targeted by the SEPs: **(a)** *E. coli* ATCC 11775, **(b)** *E. coli* AIC221, **(c)** *E. coli* AIC222, **(d)** *K. pneumoniae* ATCC 13883, **(e)** *P. aeruginosa* PA14, **(f)** methicillin-resistant *S. aureus* ATCC BAA-1556, **(g)** vancomycin-resistant *E. faecalis* ATCC 700802, and **(h)** vancomycin-resistant *E. faecium* ATCC 700221. Polymyxin B was used as positive control and buffer with DiSC_3_-5 and bacteria were used as baseline for the calculation of the relative fluorescence values.

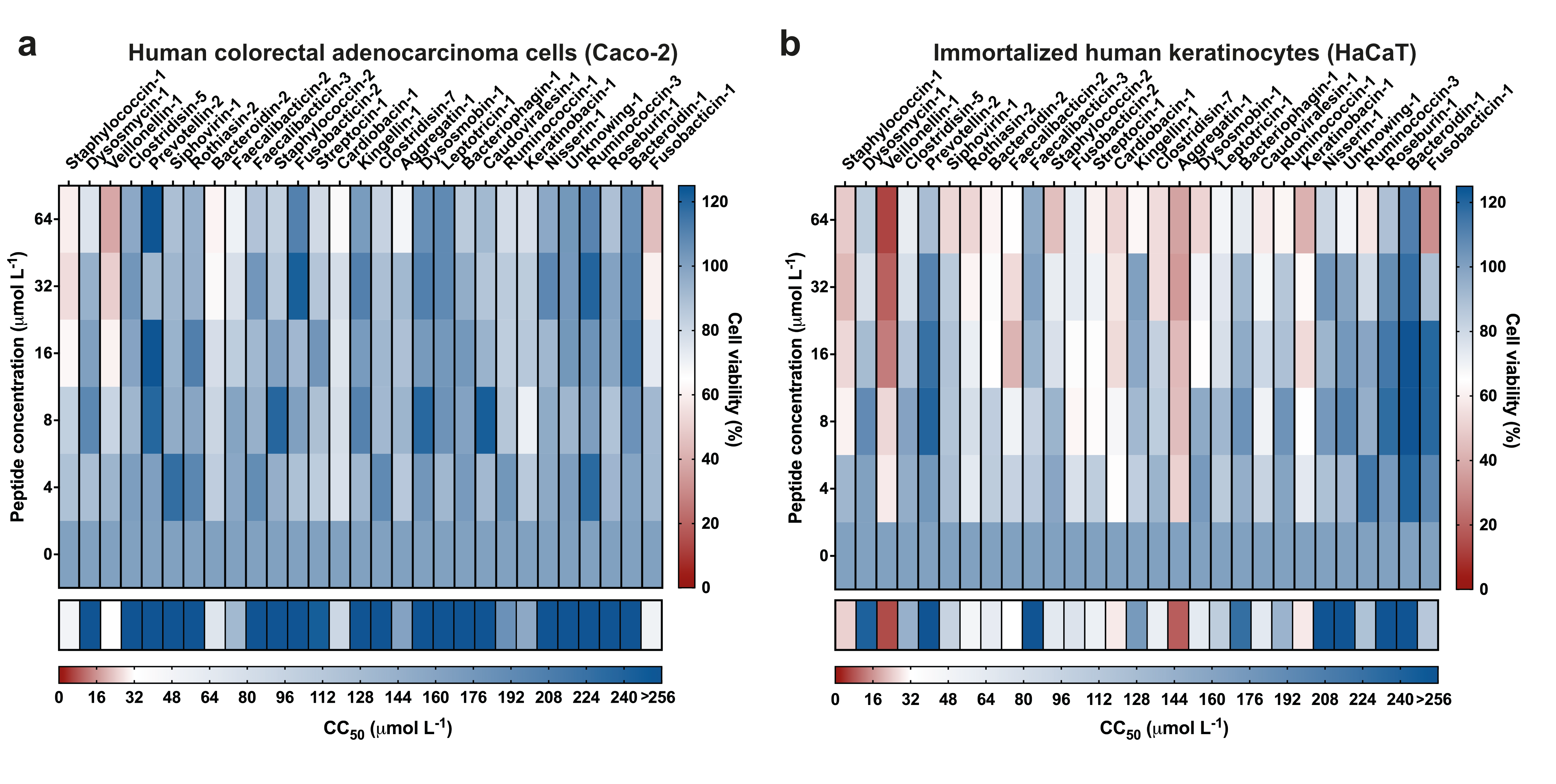

**Figure S6. Cytotoxic effects on human colorectal adenocarcinoma cells (Caco-2) and immortalized human keratinocytes (HaCaT).** Heat maps showing cell viability after 24 h of peptide treatment at concentrations ranging from 4 to 64 μmol L^-1^ against **a)** Caco-2 and **b)** HaCaT cells. Toxic (in red) and non-toxic (in blue) concentrations of the peptides are a mean of three independent replicates. The row below each of the graphs show a summary of the predicted CC_50_ concentrations (μmol L^-1^) of each peptide, i.e., peptide concentrations responsible for 50% of cell death. CC_50_ values have been predicted by interpolating the dose-response with a non-linear regression curve.
